# Amino acid and codon usage explain amino acid misincorporation rates across the tree of life

**DOI:** 10.64898/2026.02.19.706746

**Authors:** Jonas Poehls, Cedric Landerer, Katherine Daniels, Agnes Toth-Petroczy

**Affiliations:** Max Planck Institute of Molecular Cell Biology and Genetics, Pfotenhauerstraße 108, 01307 Dresden, Germany; Center for Systems Biology Dresden, Pfotenhauerstraße 108, 01307 Dresden, Germany; Cluster of Excellence Physics of Life, TU Dresden, 01062 Dresden, Germany

## Abstract

Protein translation is an error-prone process resulting in a random population of altered protein sequences in every cell. Here, we analyzed thousands of publicly available mass spectrometry datasets to detect amino acid misincorporations and quantify error rates in 14 model organisms. We find that overall error rates and the patterns of codon to amino acid error rates correlate across species. We estimate that on average 1-2% of protein molecules in a cell harbor a misincorporation, whereas this proportion can reach 10% for long proteins. Highly expressed and very long proteins have lower error rates, indicating evolutionary selection on codon usage to reduce the cost of translation errors. While both codon-anticodon mispairing and tRNA mischarging contribute to misincorporations, we estimate that ∼70% of misincorporation events are due to mispairing. The more frequent an amino acid in the proteome, the more likely it is misincorporated (r = 0.53), likely because frequent amino acids are abundant in the cell, increasing the rate of mischarging, and have abundant tRNAs, leading to increased mispairing. Overall, we find that amino acid and codon usage explain error rates. The conserved patterns of amino acid misincorporations from bacteria to humans suggest universal mechanisms driving translational fidelity.

## Introduction

The transfer of information from DNA to mRNA to protein, a fundamental process of life, is prone to errors. These errors may occur during either transcription or translation, and are collectively known as phenotypic mutations^1–3^. Protein translation in particular can deviate from the standard genetic code in various ways, for example via stop codon readthrough, ribosomal frameshifting, or single amino acid misincorporations. Common to all phenotypic mutations is that they occur stochastically, such that at a given position in the sequence, a fraction of ribosomes will make an error. This error rate depends on the sequence context^4–9^ as well as environmental conditions^6^ and the abundances of reaction partners, such as tRNAs^10–12^, amino acids^13^, or proteins that are involved in tRNA editing^14^ or translation termination or elongation^15,16^. This means that, through codon choice or the expression levels of relevant factors, organisms may control the rate of a given error. In certain cases, individual errors occur at very high rates and fulfill an adaptive function. Such so-called programmed errors include cases of ribosomal frameshifting^17–23^ or stop codon readthrough^24–28^.

In contrast, amino acid misincorporations, due to their smaller impact on the protein sequence and structure, are often considered random noise. Yet, the translation machinery, the tRNA pool and codon usage were found to be optimized for reducing the deleterious impact of harmful misincorporations^29^. While there is selection pressure to reduce translation errors, in a few cases, high rates of amino acid misincorporations are associated with adaptive functions. In *C. albicans*, CUG mistranslation from leucine to serine, which naturally occurs at a rate of 3%, generates cell surface variability, which was shown to significantly alter fungus-host interactions^30^. In mammals, ∼1% of methionines (Met) are aminoacylated to non-methionyl-tRNAs. This rate is even higher upon stress, such as oxidative stress or viral infection. It was suggested that Met-mischarging to non-methionine tRNAs may have an adaptive function to increase Met incorporation into proteins to protect cells against oxidative stress^31^. However, to assess the full impact of misincorporations, be it as deleterious noise or adaptive recoding, a proteome-wide picture of the rate of misincorporations is needed.

Detection of misincorporations is challenging, since they are generally rare and occur stochastically. Their proteome-wide detection is only possible using mass spectrometry-based proteomics (MS). In mass spectrometry, an amino acid misincorporation compared to a reference sequence will manifest as a predictable shift in the mass of a peptide, equal to the mass difference between the original and misincorporated amino acid. The identification of peptides with mass shifts using an algorithm such as ModifiComb^32^, MSFragger’s ‘open search’^33^ or MaxQuant’s ‘dependent search’^34^ allows, after appropriate processing, the quantification of misincorporations. This approach was pioneered a few years ago^35^, and has recently been employed to detect misincorporations in hundreds of datasets from *E*.*coli* and *S. cerevisiae*, revealing adaptation of the translation machinery^29^, and in *D. melanogaster* over the course of development^36^. However, often only a few sites and peptides were detected, preventing quantification of all error rates^35^. In addition, a comparative analysis across diverse organisms is still lacking.

Here, we analyzed 3,204 proteomics datasets of 14 model organisms and identified misincorporations at more than 100,000 sites. Misincorporation rates correlate between most species pairs, and we are able to identify consistently error-prone codons and amino acids. We see that amino acid and codon usage correlate with observed error rates, showing that organisms can influence translation errors via codon choice and metabolism. We find evidence of selection against translation errors in highly expressed and very long proteins. In summary, we provide a comparative overview of misincorporation patterns including misincorporation rates of individual codon-to-amino acid pairs, per-site and per-protein error rates.

## Results

### Quantification of codon-to-amino acid error rates in 14 species

To accurately measure amino acid misincorporation rates, we reanalyzed publicly available proteomics data at large scale. For 14 different model species, we collected all PRIDE^37^ datasets which were derived from a single species and acquired on an Orbitrap-family mass spectrometer, which allowed us to process all data using a single workflow.

We analyzed the proteomics data using the deTELpy pipeline (Fig 1A), which is based on the ‘open search’ algorithm^33^ implemented in the MSFragger search engine^38^. Briefly, the open search algorithm allows the identification of peptides containing arbitrary mass shifts. By processing and filtering all mass shifts observed in an MS measurement (see Methods for details), we identify amino acid substitutions with exact localization. Thus, we can determine at which position of which protein an incorrect amino acid was inserted, and by combining this with the proteins’ coding sequences, we can determine which codon was (mis-)translated as which amino acid.

**Figure 1:**
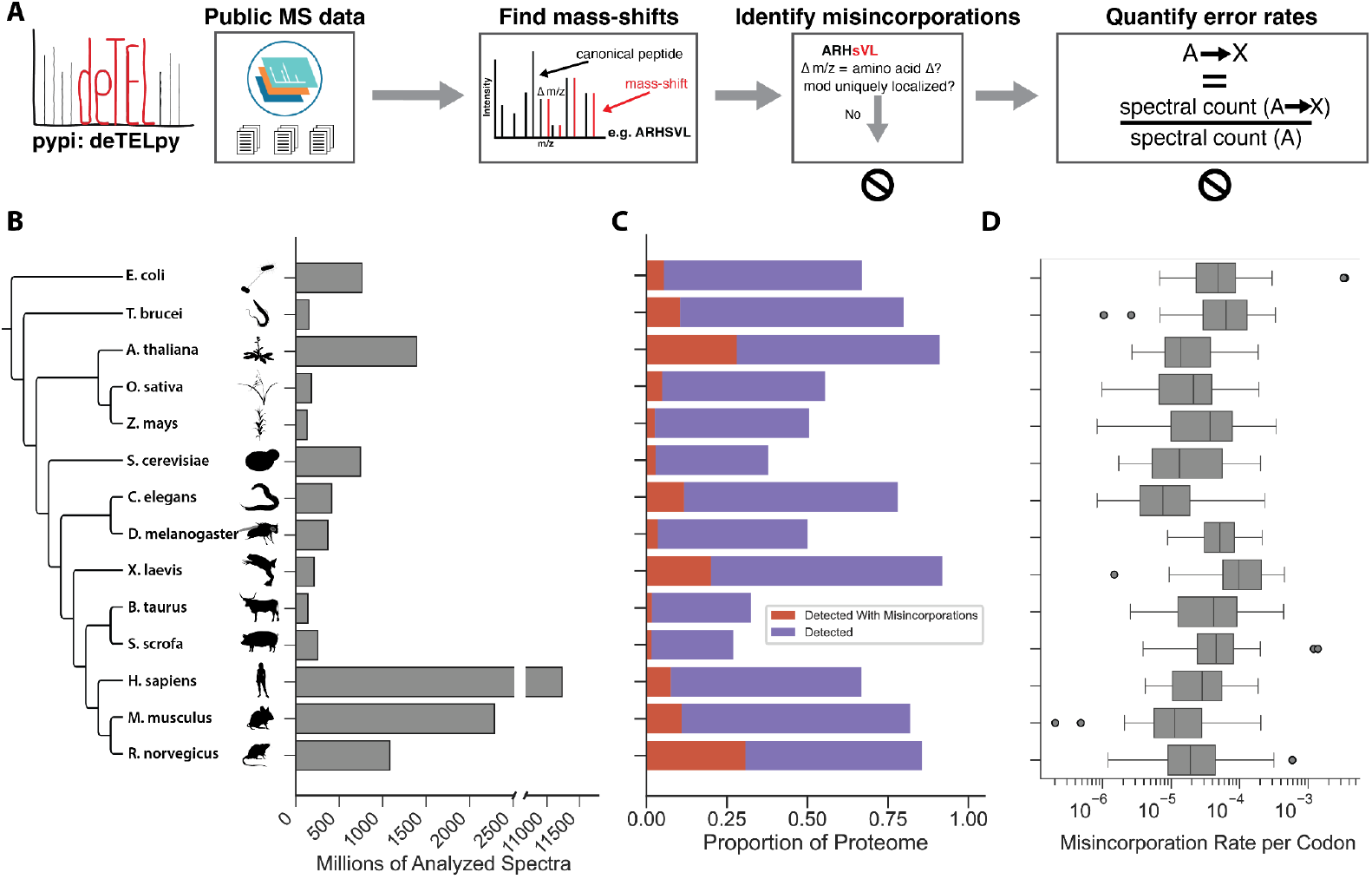
Large-scale detection of amino acid misincorporations in 14 model organisms. A. Mass spectrometry-based pipeline to quantify amino acid misincorporations using deTELpy. Error rates are computed as the ratio of the counts of codon (A) to amino acid (X) misincorporations observed across all samples divided by the codons observed (spectral counts) without an error. B. After selecting and aggregating proteomics data from PRIDE, the analyzed data for individual species ranged from ca. 100 mio. spectra for maize to more than 11 bio. spectra for humans. C. Proportion of proteins detected across all datasets, either canonical or canonical and with at least one misincorporation. D. Boxplots show the total error rates (summed over amino acids) for each codon across species. Total error rates range from less than 10^-6^ to more than 10^-3^.

To quantify the rate of a given codon-to-amino-acid misincorporation type, we do not rely on the peptides’ MS signal intensities. Instead, we make use of the fact that we observe each misincorporation type many times independently in different pairs on canonical and misincorporated peptide. To calculate the rate of a given error (given codon mistranslated as a given amino acid), we count how often the codon has been observed (number of matched spectra) both with and without a misincorporation, summed across all observed positions with this codon. Then the misincorporation rate is calculated as the number of identified spectra where the misincorporation was observed divided by the total number of spectra where the given codon was observed.

We applied the approach described above to measure the patterns of amino acid misincorporations in 14 different species encompassing bacteria, protists, plants, fungi, and animals including humans (Fig 1B, left). For each species, we processed between 17 and almost 2,000 datasets, containing 100 mio. (for maize) and more than 11 bio. (for humans) spectra (Fig 1B, left; Suppl. Table 1). From the identified misincorporations, we removed likely false positives by filtering outlier datasets, outlier positions, immunoglobulins (due to their variable region), and common laboratory contaminant proteins (Suppl Fig 1).

Using this workflow, we detected misincorporations in hundreds to thousands of proteins in each investigated species, representing between 1.6% and more than 30% of the proteome (Table 1, Fig 1C). The misincorporations occur at ca. 100,000 distinct proteome positions, ranging from ca. 1,000 for *Z. mays* to almost 45,000 for humans (Table 1, Suppl. Fig 2). Considering our strict filtering and the generally incomplete coverage of MS-based proteomics, it is likely that misincorporations occur in even more proteins and positions than we detected. Based on the detected misincorporations, we calculated total per-codon error rates as well as the rates of individual codon-to-amino-acid misincorporations types. The average error rate across all codons is within ca. half an order of magnitude for all species, between ca. 4*10^-4^ and 4.7*10^-4^ (Table 1). Per-codon total error rates are between 10^-5^ and 10^-3^ for most codons in most species, with low outliers in the 10^-6^ range and high outliers in the 10^-3^ range (Fig 1D). When looking at the mean rate across species, the rates of individual misincorporation types differ significantly, spanning five orders of magnitude from less than 10^-9^ to more than 10^-4^ (Fig 2A). We observe that some misincorporations occur at high rates for all synonymous codons, such as alanine to glutamine or proline to alanine, while other misincorporations depend strongly on the identity of the codon, such as arginine to tryptophan, which occurs at high rates for CGG and AGG codons, but not their synonyms (Fig 2A).

**Table 1:** Observed misincorporations and misincorporation rates for 14 model organisms.

| Species | Proteins with misincorporations | % Proteome with misincorporations | Misincorporated positions | Log10 error rate |
| --- | --- | --- | --- | --- |
| *E. coli* | 1,177 | 30.7 | 11,062 | -4.046 |
| *T. brucei* | 943 | 11.0 | 1,837 | -4.046 |
| *A. thaliana* | 2,085 | 7.6 | 7,917 | -4.523 |
| *O. sativa* | 686 | 1.6 | 1,380 | -4.523 |
| *Z. mays* | 698 | 1.8 | 1,008 | -4.398 |
| *S. cerevisiae* | 1,182 | 20.1 | 5,876 | -4.523 |
| *C. elegans* | 705 | 3.6 | 2,283 | -4.699 |
| *D. melanogaster* | 1,622 | 11.7 | 4,145 | -4.222 |
| *X. laevis* | 1,013 | 2.9 | 2,005 | -3.959 |
| *B. taurus* | 618 | 2.6 | 1,578 | -4.222 |
| *S. scrofa* | 1,114 | 4.9 | 2,761 | -4.155 |
| *H. sapiens* | 5,776 | 28.1 | 44,661 | -4.398 |
| *M. musculus* | 2,298 | 10.5 | 9,737 | -4.699 |
| *R. norvegicus* | 1,227 | 5.4 | 3,974 | -4.699 |

**Figure 2:**
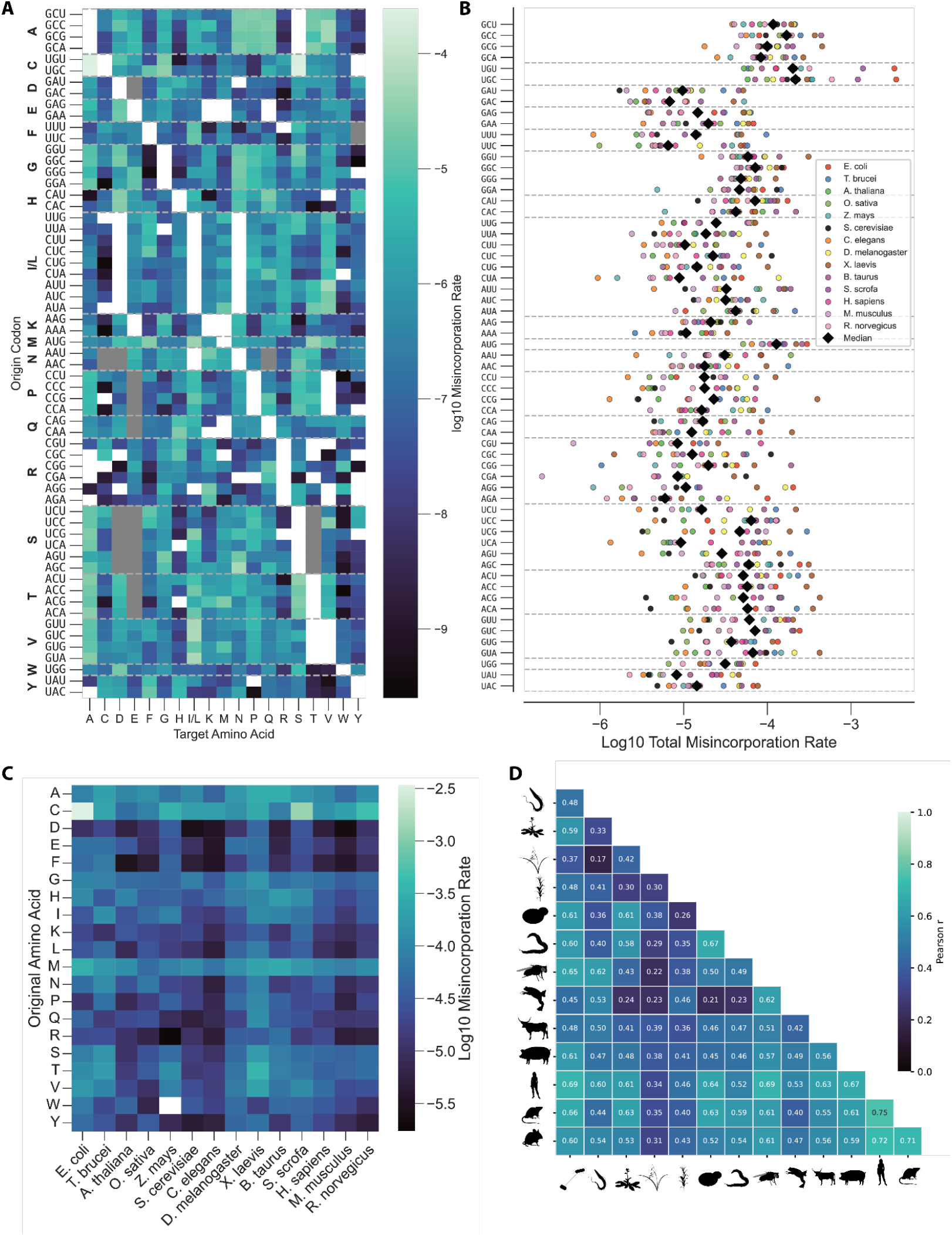
Misincorporation patterns correlate across species. A. Mean codon-to-amino acid error rates across species. Codon-to-amino acid misincorporation rates averaged across all 14 species. Each rate is calculated as the number of identified spectra where the misincorporation was observed divided by the total number of spectra where the given codon was observed. White fields indicate misincorporations that were never observed. Gray fields indicate misincorporations that cannot be observed because they are masked by post-translational modifications or artifacts with the same mass and thus undetectable. Individual codon-to-amino acid error rates range from less than 10^-9^ to more than 10^-4^. B. Total misincorporation rates per codon across species. Shown are the total misincorporation rates (summed across destination amino acids) for each codon in each species. The median is indicated by a black diamond. Individual rates range from less than 10^-6^ to more than 10^-3^. C. Total misincorporation rates by origin amino acid and species. Total misincorporation rates were summarized at the level of the origin amino acid by combining total observations and misincorporations of all codons coding for a given amino acid. Alanine, cysteine, and methionine are overall the most error-prone amino acids, while phenylalanine and aspartate show the lowest error rates. D. Pearson correlation coefficients of codon-to amino acid error rates between species. Only the rates of misincorporation types observed in both species were compared.

When comparing individual misincorporation rates between species, the error rates of some codons, such as those coding for lysine or glycine, are relatively similar between species, while others, for example CCG or UUU, differ by up to two orders of magnitude between individual species (Fig 2B).

By combining synonymous codons, we can calculate the total misincorporation rate of each amino acid and compare these between species (Fig 2C). The most error-prone amino acids are alanine, cysteine, and methionine, while phenylalanine and aspartate tend to be the least error-prone.

The misincorporations patterns of the species we examined are similar to each other: The Pearson correlation coefficients of codon-to-amino-acid error rates between species pairs are mostly greater than 0.4. The highest correlation is seen between human and rat (r = 0.75), and all five mammals included in our data have a correlation coefficient of 0.5 or higher with each other. *X. laevis, O. sativa*, and *Z. mays* show weaker correlation, including with each other (Fig 2D).

The misincorporations patterns allow us to identify factors explaining the observed error rates, and to investigate the degree of conservation of errors between species.

### Protein properties and codon choice explain patterns of misincorporation errors

After observing that the rates of specific error types differ by several orders of magnitude, we wanted to identify factors that can explain these differences. It is generally assumed that highly expressed proteins are subject to stronger selection on their codon choice. Since there is some evidence that preferred codons are translated not only more efficiently, but also more accurately, we tested whether there is a relationship between the abundance of a protein and its per-protein error rate. We approximated protein abundance by the normalized spectral abundance factor (NSAF), which is equal to the total spectral count divided by the protein’s length. Per-protein error rates are equal to the spectral count of peptides with misincorporations mapping to the protein divided by the protein’s total spectral count.

For all species observed, except *E. coli*, we see a significant negative correlation between NSAF and per-protein error rate, with Pearson’s correlation coefficients between -0.11 and -0.68 (Suppl Fig 4), showing that highly expressed proteins contain fewer misincorporations.

Amino acid misincorporation is based on binding events, the probabilities of which are dependent on the reactant’s concentrations. We thus hypothesized a relationship between the cellular abundance of an amino acid and its tRNAs and the rate at which it is incorrectly incorporated. We observed a moderately strong positive correlation between the frequency of an amino acid in the organism’s proteome and the total rate of errors where the amino acid is incorporated (Pearson’s r = 0.53, p = 2.62*10^-19^). This relationship holds both for combined data from all species (Fig 2B) and for 12 out of 14 individual species (Suppl. Fig. 5), indicating that more frequently used amino acids are more likely to be misincorporated.

We observed significant differences between the error rates of synonymous codons (Fig 2B). We were interested to see whether these tendencies are conserved between species, i.e. whether within a codon group there are codons that are, across species, consistently more error-prone than synonymous codons. To quantify this, we considered each group of synonymous codons separately and, for each species, ranked them according to their total error rate (i.e. rank = 1 means the codon has the highest error rate among synonymous codons). We then calculated each codon’s mean rank across organisms. We compared this to the expected distribution of mean ranks if ranks were assigned randomly and calculated empirical p-values as a measure of how extreme the actual ranks are compared to the naive expectation (see Methods for details). When applying a threshold of p = 0.05, nine codons are significantly different from the expectation; four codons tend to have lower error rates than their synonyms: AAA (Lys), UCA (Ser), GUG (Val), and UAU (Tyr); while five codons tend to have higher error rates: GCC (Ala), GGC (Gly), AAG (Lys), CGG (Arg), and UAC (Tyr) (Fig 3C).

**Figure 3:**
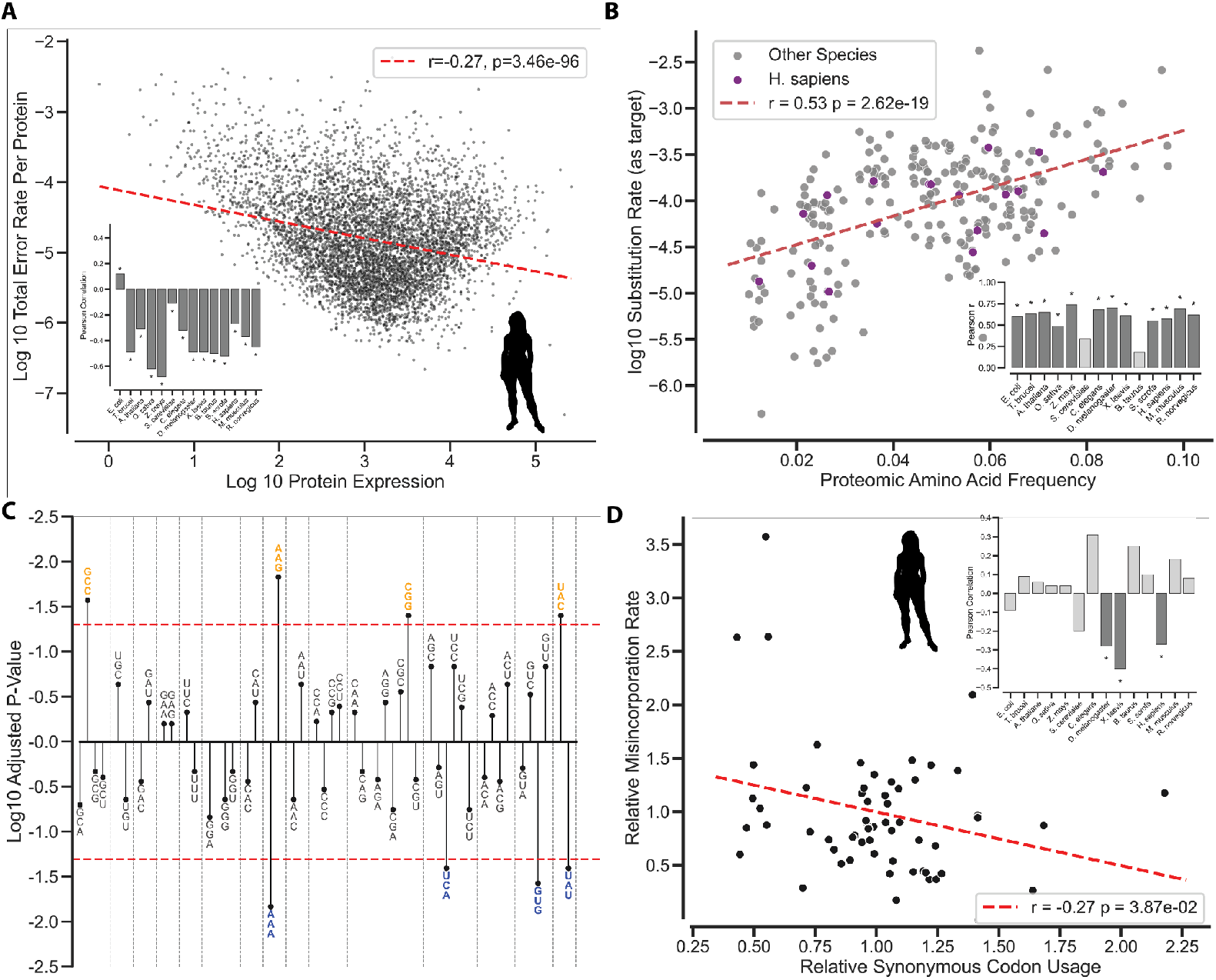
Amino acid and codon usage correlate with misincorporation error rates. A. Per-protein error rates decrease with protein expression. Shown are total error rates versus normalized spectral abundance factor (NSAF, spectral count divided by protein length) for all human proteins with a non-zero error rate. Per-protein error rate and NSAF are significantly negatively correlated (r: Pearson correlation coefficient, p-value: two-sided Wald test) for all species analyzed, except *E. coli* (Suppl Fig 4). B. Amino acids appearing more frequently in the proteome are more frequently misincorporated. Scatter plot shows that amino acids frequency (proportion of positions in the proteome with this amino acid) versus their total misincorporation rate (sum across all misincorporation types where a given amino acid is misincorporated) for all species are significantly positively correlated (Pearson r = 0.53, p = 2.62*10^-19^, two-sided Wald test). This trend holds at the level of individual species for 12 out of 14 species (Suppl. Fig 5). Inset: Pearson correlation coefficients for individual species. * indicates p <= 0.05. C. Relative error tendencies of synonymous codons are partly conserved between species. Within each species, synonymous codons were ranked by error rate. Empirical P-values were calculated by comparing codon’s mean rank across species to the distribution expected for random assortment of ranks (Suppl Fig 6, see Methods for details) and adjusted for multiple testing by Benjamini-Hochberg correction. Low p-values indicate consistently high or consistently low error rate compared to synonyms across species. Codons without synonyms (AUG, UGG) are not shown. Dotted red line corresponds to adjusted p = 0.05. D. Scatterplot of relative synonymous codon usage (RSCU) and relative misincorporation rate (RMR) of each codon for humans. RMR is a codon’s total error rate divided by the mean total error rate of its synonyms. Inset: P-values of two-sided Wald’s test of RSCU and RMR for all species. RSCU and RMR are significantly negatively correlated in humans, *D. melanogaster*, and *X. laevis*.

Since synonymous codons differ significantly in their error rates, we tested whether there is a relationship between relative synonymous codon usage (RSCU) and error rate. We normalized each codon’s total error rate by the mean error rate of its synonyms with each species to obtain the relative misincorporation rate in order to isolate the effect of the codon choice. For 3 of the 14 species (*H. sapiens, D. melanogaster, X. laevis*), we see a significant negative correlation between RSCU and the relative misincorporation rate, although even in these the effect is limited, with Pearson’s r between -0.27 and -0.4 (Fig 3D, Suppl Fig 7).

In summary, we find that amino acid frequencies and codon usage help explain the observed misincorporation rates. The relationship of error rate with protein abundance and codon usage points towards selection pressure on the coding sequence to mitigate translation errors.

### Long proteins are encoded by less error-prone codons

Our dataset of codon-to-amino acid error rates enabled us to estimate the probability that a given protein molecule will contain at least one error. The probability that the protein will be produced with at least one misincorporation is the cumulative product of the error-free probability (1 - total misincorporation rate) of each codon in the protein’s sequence, subtracted from 1, hereafter referred to as the error proportion (see Methods for details). The error proportion generally lies between 0 and 10% for most proteins, with a peak around 1-2% for most species (Fig 4A, Suppl. Fig 8). The median across the whole proteome lies between ca. 0.5% for *C. elegans* and more than 5% for *X. laevis* (Fig 4B).

**Figure 4:**
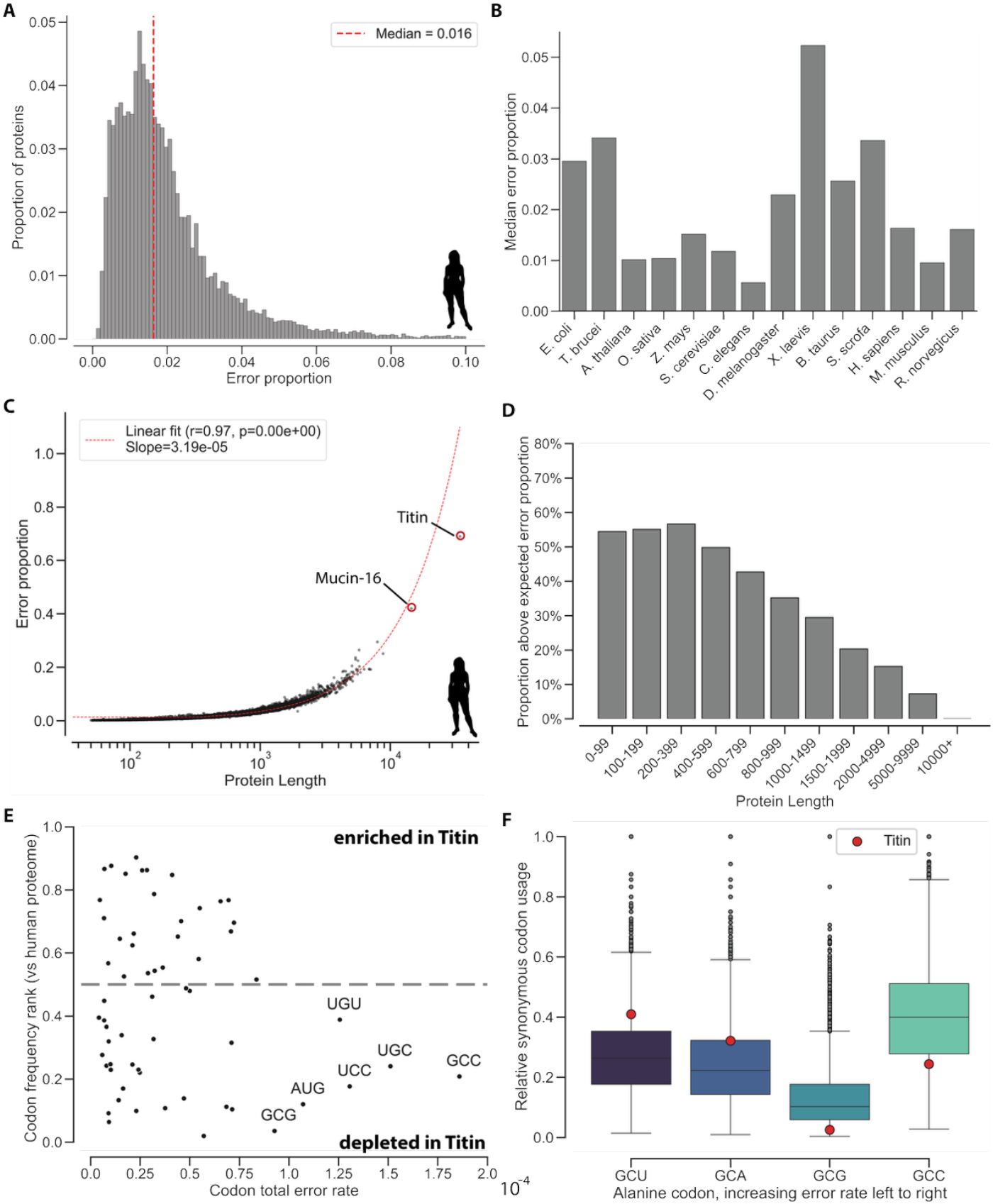
Long proteins are encoded by less error-prone codons. A. Histogram of expected error proportion of human proteins. By combining the total misincorporation rate of each codon along each protein’s coding sequence, we obtain the protein-specific probability to contain at least one misincorporation, termed error proportion. For humans, the median error proportion is 1.6%, and the highest values are around 10%. B. Median expected error proportion across proteome per species. Median values are mostly between 0.5 and 3%, with *X. laevis* being the outlier at more than 5% due to the elevated error rates measured in this species. C. Relationship of protein length and expected error proportion for human proteins. r: Pearson correlation coefficient. P-value calculated using two-sided Wald test. The slope represents the average increase in error proportion with each codon. Note that the line represents a linear regression, but appears exponential due to the logarithmic x-axis. The two human proteins with >10,000 codons, mucin-16 and titin, lie below the regression line, indicating they have a lower length-normalized error proportion than the average protein. D. Proportion of proteins with a higher-than-expected expected error proportion by length. For each species, the regression line of protein length and error proportion gives the expected error proportion for a given protein length. The bar chart shows the proportion of proteins in each length bin that lie above this regression, for all species combined. For all length bins above 600 residues, the minority of proteins are above the expectation, and the proportion becomes smaller with each increasing length bin, indicating that longer proteins use less error-prone codons. E. Human titin is depleted in error-prone codons. Plotted is the frequency percentile rank (proportion of human proteins with lower frequency of codon than titin) and total misincorporation rate in humans for each codon. The six codons with the highest error rates are less frequent in titin than in the majority of human proteins. F. Human titin has alanine codon usage adapted for lower misincorporation rates. Boxplot shows the relative synonymous codon usage of alanine codons for human proteins. Codons are ordered by total misincorporation rate increasing left to right. Titin uses the two least error-prone codons more and the two most error-prone codons less than the average protein.

Since each codon provides an opportunity for a misincorporation, the error proportion naturally scales with the protein’s length. This correlation is very strong in all species considered here, with correlation coefficients between 0.89 and 0.99 (e.g. humans: Pearson’s r = 0.98, Fig 4C, Suppl. Fig 9). Note that the slope of the regression line is equal to the average per-codon increase in error proportion.

However, the relationship between length and error proportion is not perfect. Indeed, we noticed that extremely long proteins, such as titin and mucin-16 in humans, tend to lie below the regression line. Titin, for example, has an error proportion of 69%, but if it would lie on the regression line, this value would be close to 100% (Fig 4C). Thus, if titin would have a similar per-codon error rate as the average protein, there would hardly be a titin molecule matching the gene sequence.

We tested this trend more systematically by comparing the error proportion of each protein to the naïve expectation, which is simply the average per-codon increase in error proportion multiplied by the protein length. Indeed, we find that the longer the protein, the more likely it is to have a lower-than expected error proportion. When binning the proteins by length, starting from the 600-799 bin, the majority of long proteins have a lower-than-expected error proportion, and this proportion gets larger with each increasing length bin, until almost all proteins above 10,000 amino acids are below the expectation (Fig 4D). Since this analysis is based only on the codons’ global error rates, the only explanation for this is that long proteins use less error-prone codons.

We investigated the different codon usage of long proteins in more detail, using human titin as an example. For each codon, we calculated the proportion of human proteins that use a codon less frequently than titin. A value below 0.5 thus indicates depletion of the codon in titin compared to the proteome. The six codons with the highest total error rate in humans are depleted in titin. This includes the codons for cysteine and methionine, the most error-prone serine codon UCC, and the two most error-prone alanine codons, GCG and GCC (Fig 4E). While in the case of cysteine and methionine this might simply indicate a lower need of these amino acids in the protein, for alanine and serine, this hints at different usage of synonymous codons. When looking at the relative usage of alanine codons, we can confirm that titin uses the two less error-prone codons, GCU and GCA, more than most human proteins, and uses GCG and GCC less (Fig 4F). This indicates that the codon usage of titin is adapted to avoid amino acid misincorporations.

Thus, long proteins tend to use less error-prone codons than the average protein. Since both the production and the degradation of such large proteins is extremely costly, this could be an evolutionary adaptation to reduce the proportion of error-containing molecules and thus reduce the cost of misincorporations.

### Misincorporation errors arise mostly from non-Watson-Crick pairs involving G or U and tRNA mischarging

There are two possible mechanisms that can lead to the amino acid misincorporations that we observe: The charging of a tRNA with an amino acid not matching its anticodon (“mischarging”) and the binding of an anticodon to a non-cognate codon (“mispairing”). While mispairing and mischarging errors are indistinguishable based on the protein sequence alone, we can infer the error type by assuming that the more basepair mismatches between codon and anticodon would be needed, the less likely a mispairing error is. We therefore inferred the codon-anticodon mismatches required for all observed misincorporations, based on the differences between the codon and the amino acid’s most similar anticodon (see Methods for details).

Most misincorporations we observe would require a mismatch in the 1st codon position, the 2nd position, or both (Fig 5A). Misincorporations requiring a single mismatch are the large majority, followed by two, and less than 10% of misincorporations would require three mismatches. Misincorporations requiring mismatches in the 3rd position are rarer, which is likely due to the structure of the genetic code, where many third-position mismatches would not lead to a change in the amino acid. We nevertheless take the fact that almost 40% of observed misincorporations require at least two mismatches as an indication that we mostly observe genuine translation errors, since two consecutive genetic or transcriptional errors should be very rare.

**Figure 5:**
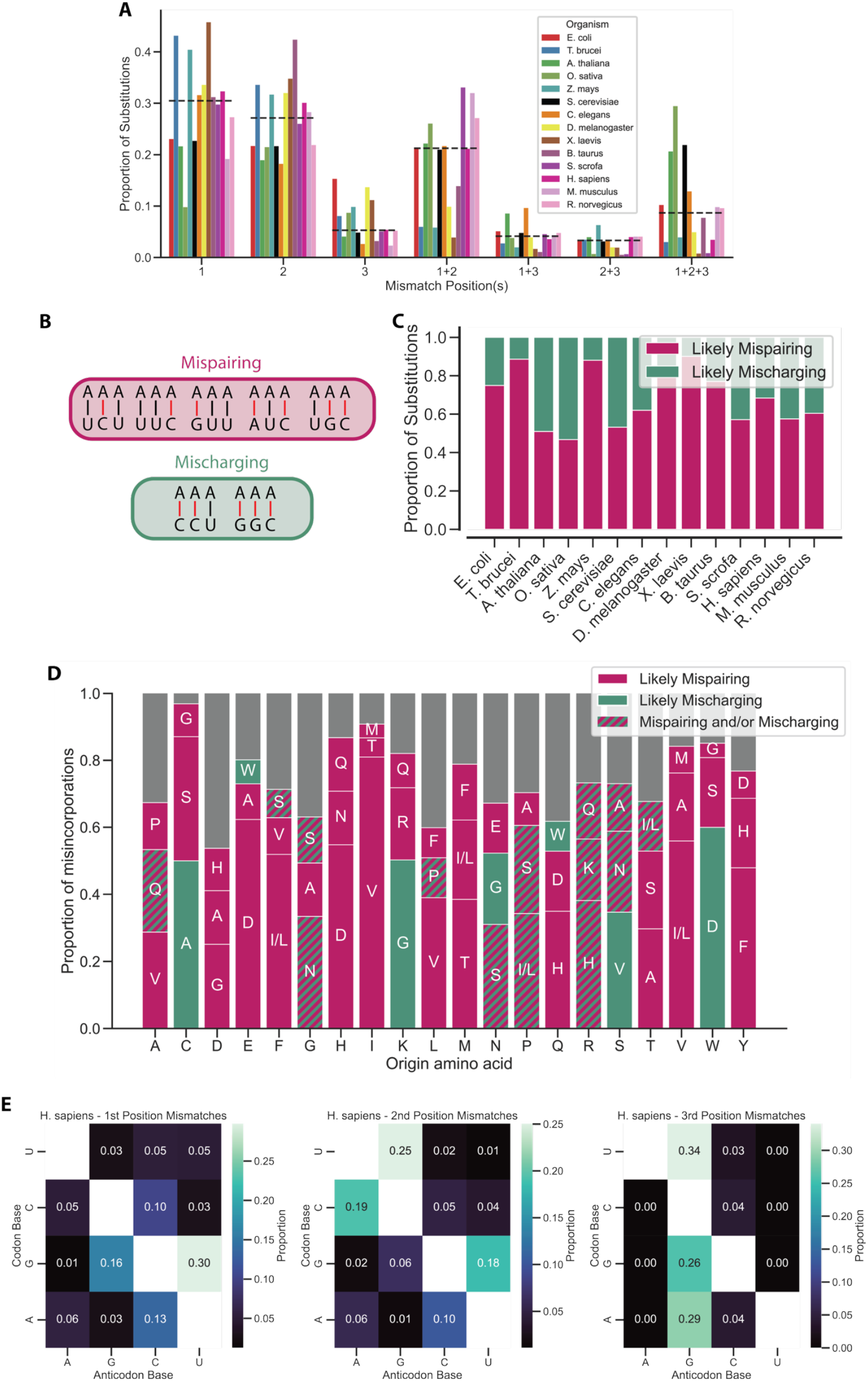
Majority of misincorporation errors arise from non-Watson Crick pairs conserved across species. A. Amino acid misincorporations at the translation level might be due to either mischarging of the tRNA, resulting in a mismatch between the anticodon and the amino acid, or due to mispairing of the codon with a non-cognate anticodon. We classify all misincorporations requiring mismatches at the 1st and 2nd position as likely mischarging, and all others as likely mispairing. Black dotted line represents median across species. B. Majority of misincorporations would require single mismatch at 1st or 2nd position. Shown is the distribution of the position(s) of the mismatches between a codon and the closest-matching anticodon of the misincorporated amino acid across all observed misincorporations. Ambiguous cases where more than one anticodon is equally close were excluded from this analysis. C. Most misincorporations are due to codon-anticodon mispairing. Based on theoretically required mismatches, misincorporations were classified as either likely mispairing (near-cognate codon-anticodon binding) or likely mischarging (charging of tRNA with wrong amino acid). For all species except *O. sativa*, the majority of misincorporations are likely mispairing. D. Distribution of misincorporated amino acids is highly specific to the encoded amino acid. Bar chart shows the proportions of the three most commonly misincorporated amino acids based on the originally encoded amino acid. Grey bars show the summed proportion of the remaining amino acids. Colors indicate the inferred misincorporation type. Hatched bars indicate that either mispairing or mischarging is possible, either because the type depends on the codon, or because the type cannot be inferred (‘Ambiguous’, see Methods for details). E. Proportions of inferred nucleotide mismatches for likely mispairing misincorporations by codon positions, for humans (see Suppl Fig 10 for all species). Proportions are relative to all mispairing misincorporations with a mismatch at the respective position. In 1st and 2nd position, most misincorporations are due to same-nucleotide and pyrimidine-purine mismatches, particularly G-U matches. In the 3rd position, almost all misincorporations involve a G in the anticodon.

We then classified the observed misincorporations as either likely mispairing (single mismatch or two mismatches involving the 3rd position) or likely mischarging (mismatches at 1st and 2nd position) (Fig 5B). With this classification, the majority of errors are likely due to mispairing for all species except rice. The proportion of mispairing errors ranges from 46% (rice) to 88% (*Xenopus*) and is 68% for humans (Fig 5C), thus most errors are the result of tRNA mispairing. This trend is also clear when considering misincorporations from the perspective of the originally encoded amino acid: For most amino acids, the most common misincorporations are likely due to mispairing, however there are some very common mischarging misincorporations as well, such as C->A, K->G, or W->D (Fig 5D). These might be due to common mischarging by alanyl-tRNA synthetase which is known to mischarge alanine onto cysteine tRNA in human but not in E.coli^39^.

In addition, the misincorporated residues are highly specific to the originally encoded amino acid: For all ‘‘origin amino acids, the three most common ‘targets’ account for more than 50% of misincorporations, and in some cases a single target accounts for more than 50% of errors, such as W->D (60%), E->D (62%), or I->V (81%) (Fig 5D).

We then investigated which nucleotide mispairings are responsible for the occurrence of the observed mispairing errors. For each of the three codon positions, we calculated the proportion of non-Watson-Crick pairings among misincorporations that have a mismatch in this position.

In humans, the most common mismatch types are G-U in the 1st position; U-G, G-U, and C-A in the 2nd position; and any mismatch involving a G in the anticodon in the 3rd position (Fig 5E). Notably, the probability of a mismatch between A and B seems to depend on whether A is part of the mRNA and B part of the tRNA or vice versa. This is likely due to two reasons: One, the ribosome’s decoding center interacts with the codon-anticodon pair, leading to thermodynamics different from solution. Two, the anticodon bases might carry modifications, influencing their binding affinities^40,41^. The high proportion of mispairings between U and G is likely because G/U pairings are able to adopt a Watson-Crick–like conformation in the ribosome’s context, and are thus tolerated similarly to Watson-Crick pairs^42^. We conclude that a subset of mismatch types, predominantly pairings involving G and U, are responsible for the majority of mispairing misincorporations.

Mismatch patterns in 1st and 2nd position are moderately to strongly conserved between most species, with Pearson’s r above 0.3 in most cases, and strongly correlated in the 3rd position, although this is likely inflated by most of the values being close to 0 (Suppl. Fig. 11).

We conclude from this that both codon-anticodon mispairing and tRNA mischarging play a role in determining misincorporation rates. Most misincorporations stem from a small subset of nucleotide mispairings and tRNA mischarging events. Patterns of nucleotide mismatching are partially conserved between species, especially among the closely related mammals.

## Discussion

Amino acid misincorporations have long been known to occur at rates of 10^-4^ or higher^43–45^ based on measurements of individual sites or proteins. More recent proteome-wide studies have focused on either a single species^36,46,47^ or at most compared two species^29,35,48^. Here, we present the first comparison of misincorporation patterns in a range of model organisms. Our analysis is based on the reanalysis of publicly available proteomics data at large scale. This aggregation has two main advantages: One, due to the bias for highly abundant proteins in mass spectrometry^49^, misincorporations, which are generally rare, are difficult to detect. Combining large amounts of data allows us to detect many individual misincorporations and thus to estimate misincorporation rates more accurately and for a larger number of misincorporation types. Two, estimation of error rates requires us to compare the amounts of mistranslated and canonical protein products and thus peptides. This is inherently difficult in MS-based proteomics, as the signal intensities of different peptides, even with a difference of a single amino acid, are generally not directly comparable, especially if peak areas are used^50,51^. We partially address this issue by not comparing individual pairs of mistranslated and canonical peptides (as done before^35,47,48^), but rather combining all observations of a given misincorporation type and comparing total amounts of misincorporated and canonical products.

Across species and misincorporation types, the error rates we estimate span at least five orders of magnitude from below 10^-9^ to more than 10^-4^. We observe that the identity of the codon and of the (mis-)incorporated amino acid seem to be the dominant factors determining the misincorporation rate, rather than the species or the protein. This is supported by the fact that error rates are moderately to strongly correlated between species, and the observation that the correlation between amino acid frequency and misincorporation rate is stronger than that between protein abundance and error rate for most species (compare Suppl. Figs. 4 and 5).

We identify several factors that correlate with error rates. The negative correlation between a protein’s abundance and its error rate we observe might point to a stronger selection against misincorporations in more highly expressed proteins. This point is related to our observation that extremely long proteins tend to use less error-prone codons. Assuming that protein copies with misincorporations would have reduced function or even be harmful, error-prone codons in highly expressed or very long proteins would incur larger cost to the cell than in less abundant or shorter proteins and thus more likely to be subject to selection. Based on this, we would expect misincorporation rates to correlate with measures of codon optimality such as RSCU. While we do see correlations between error rate and RSCU, the trend is only statistically significant for a few of the species we investigated.

The observation that amino acids appearing more frequently in the proteome are misincorporated more often might be explained by a combination of basic chemical principles and cellular optimization of translation: Presumably, frequently used amino acids such as alanine or serine, as well as their tRNAs, are more abundant in the cell. There is evidence that tRNAs compete for binding to the ribosome and that misincorporation patterns are partially explained by tRNA abundances^10,29^, and similarly, tRNA mischarging rates might depend on amino acid abundances^13^. Indeed, amino acid concentrations in the cells influence misincorporations: Arginine deprivation enriches lung cancer proteomes with cysteine by inducing arginine-to-cysteine misincorporations^52^ and tryptophane depletion leads to Trp-to-Phe misincorporations^13^ in cancer cells.

Based on the differences between a codon and the anticodons encoding the misincorporated amino acid, we infer whether misincorporations are due to mispairing or tRNA mischarging. A limitation of this is that while misincorporations that would require improbable mismatches as likely due to mischarging, any misincorporation which we label as likely mispairing could in principle also be due to mischarging. Similar to other studies^35,36^, this implicitly assumes that mispairing errors are more likely and should be the default inference unless there is evidence for mischarging. We are not aware of any direct comparison of mischarging and mispairing rates, which would test this assumption.

An important limitation of investigating translation errors using the approach employed by us and others is that it does not measure the amount of errors made, but at best the amount of error-containing products present. Should the lifespan of erroneous proteins differ from that of canonical ones, e.g. through selective degradation, this would bias the apparent error rates. In addition, while aggregating many studies allows greater coverage and depth, the various treatments, enrichments, and sample preparations require us to consider a large number of possible PTMs. This results in 11 amino acid misincorporations being undetectable with our pipeline (Suppl. Table 2).

It is unclear to what degree the misincorporations observed here are functional or adaptive. In a few cases, individual misincorporations have been shown to be adaptive, the most famous example being methionine misincorporations under oxidative stress^31,53,54^. In addition, there are both theoretical^55^ and experimental^3^ results showing that misincorporations can facilitate the evolution of proteins. We consider it likely that most misincorporations simply reflect the trade-off between speed and accuracy inherent to translation^56,57^: While more accurate translation is possible, it would reduce translation efficiency, both theoretically^58^ and (at least in *E. coli*) experimentally^59^. In *E. coli* and *S. cerevisiae*, most misincorporations seem to be evolutionarily neutral at the observed rates, indicating that there is little pressure for a further increase in translation accuracy^29^.

In summary, we provide evidence for the widespread existence of mistranslated proteins in a range of model organisms. We provide a comparative quantification to facilitate further investigation into the evolutionary effect, molecular determinants, and possible functional roles of misincorporations.

## Methods

### Dataset selection and curation

For each species, the PRIDE API was queried to select datasets tagged exclusively with that species, an Orbitrap-family instrument, and no tags relating to stable isotope labeling by amino acids in cell culture (SILAC). For selected datasets, all.raw files were downloaded via the API. After processing (see below), raw files with a clustering of mass shifts indicative of SILAC (mass shifts of +4, +6, +8, or +10 exceeding 10% of all mass shifts) were removed. Datasets without any detected misincorporations were removed from the analysis.

### Identification and quantification of amino acid misincorporations

For each selected dataset, all raw files were then processed using a slightly modified version of the deTELpy proteomics pipeline (see ref. ^29^ for a detailed description). Briefly, the raw files were analyzed using an ‘open search’ approach^33^ based on the proteomics software suite Fragpipe.

Peptide-spectrum matches (PSMs) produced by MSFragger^38^ were post-processed by Crystal-C^60^ to reduce common false positives in open searches, PeptideProphet^61^ to rescore PSM, ProteinProphet^62^ to infer proteins, and Philosopher^63^ to filter results to a false-discovery rate (FDR) of 1%. The exact commands for each individual tool were extracted using the dry-run option of Fragpipe, and the tools were then combined in a custom pipeline suitable for parallel processing in a high-performance computing (HPC) environment.

The FDR-filtered PSMs produced by the pipeline were filtered to only include protein-unique peptides. Peptides with an absolute mass shift of >5mDa were considered modified. Modified peptides were filtered according to two criteria: The unmodified peptide needed to have been identified in the same measurement (raw file), and the position of the mass shift had to be unambiguously localized by MSFragger. In addition, we compared the calculated mass shift to the masses of a list of post-translational modifications (PTMs) and proteomics artifacts and excluded all PSMs with a mass shift within 5ppm of any of these PTMs or artifacts. This results in certain misincorporations being virtually undetectable in our pipeline (Suppl. Table 2). All remaining modified peptides where the mass shift matches the mass difference between two amino acid residues were marked as an amino acid misincorporation. Based on the coding sequence of the protein, the codon, the original, and the target amino acid were annotated. Isoleucine and leucine were treated as equivalent.

For the calculation of error rates, for each organism, the spectral count of each position in the proteome was calculated as the sum of PSMs including this position. By mapping each protein position to its codon, the number of observations of each codon was calculated as the sum of spectral counts of all positions with that codon. For a given misincorporation from codon A to amino acid X, the error rate is then the number of A-to-X misincorporations divided by the total spectral count of A.

Per-protein error rates were calculated as the number of PSMs with misincorporations mapping to a protein divided by the total number of PSMs mapping to the protein. Per-dataset error rates were calculated as the ratio of a dataset’s number misincorporation PSMs and the dataset’s total PSM count. Per-position error rates were calculated by dividing the number of PSMs with misincorporations covering the position by the total number of PSMs covering the position.

### Filtering of contaminants and outliers

After identifying misincorporations and calculating preliminary error rates, misincorporations were filtered in four different ways: Contaminant proteins, immunoglobulins, outlier positions, and outlier datasets. All PSMs mapping to contaminant proteins (contaminant list used by Fragpipe) were removed. Immunoglobulins were identified as all proteins with the gene ontology term ‘immunoglobulin complex’ (GO:0019814) in their Uniprot annotation. All PSMs mapping to immunoglobulins were removed. Outlier positions were identified based on each organism’s distribution of per-position error rates: All positions with an error rate exceeding the 99th percentile of non-zero per-position error rates (after removing contaminants and immunoglobulins) were marked as outliers. Outlier positions were ignored for the purpose of calculating final error rates. Outlier datasets were similarly identified: For each dataset, the mean per-protein error rate after all previous filtering steps was calculated. All datasets with a mean error rate of more than three times the interquartile range of the organism’s datasets were marked as outliers and removed from the analysis.

PSMs (canonical and misincorporation) remaining after all filtering steps were used to recalculate final error rates at the codon, residue position, protein, and dataset levels.

### Synonymous codon error rate ranks

Within each species, codons were ranked according to their total error rate relative to their synonyms (e.g. for a six-codon amino acid, codons were assigned ranks 1 through 6 in decreasing order of their error rate). The mean rank across species was then calculated for each codon. For each ‘box size’ (number of synonyms 2, 4, 6), a null expectation for the mean rank was generated by randomly sampling ranks and taking the mean 10,000 times. For each codon, an empirical p-value was calculated as the proportion of randomly generated mean ranks of the corresponding box size that was smaller than the codon’s mean rank. P-values across codons were then adjusted for multiple testing using the Benjamini-Hochberg method^64^.

### Estimation of error proportion

To calculate the expected proportion of error-containing molecules, for each protein the error-free probability 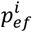 of each codon i in its coding sequence was calculated as 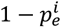, where 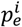 is the codon’s total misincorporation rate, summed across codons. The error proportion is then:

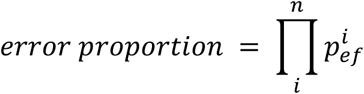

where n is the protein’s number of codons.

### Classification of mispairing vs. mischarging misincorporations

We classify a misincorporation as ‘likely mischarging’ if it would require a mismatch at both the first and second codon position (including mismatches at all three positions).

To infer the required mismatches for each observed misincorporation, the ‘origin’ codon was determined based on the protein’s coding sequence. Then, from all anticodons of the ‘destination’ amino acid present in the given species (based on the tRNA genes present in the species’ genome), the anticodon(s) with the lowest number of mismatches with the origin codons is selected as the ‘closest anticodon’.

If there is a unique closest anticodon, the position(s) and base pairs of the mismatches are annotated and the misincorporation is classified accordingly as either mispairing or mischarging. If there is more than one anticodon with the same number of mismatches, it is checked if all of them have the same mismatch position(s). If yes, the misincorporation is annotated with these positions and classified. If the anticodons have different mismatch positions, it is checked whether they would all be classified in the same way. If yes, the misincorporation is classified accordingly, otherwise it is marked as ‘Ambiguous’. For the visualization of the misincorporation types (Fig 5C), Ambiguous cases were assigned randomly to the two classes. For the analysis of mismatch positions and basepair mismatches, misincorporations without mismatch positions and basepair mismatch information, respectively, were excluded.

### Statistical analysis

For the calculation of all correlation coefficients involving error rates, the decadic logarithm of error rates was used.

All data processing and statistical analysis was done in Python 3.9 or 3.10.^65^ and Jupyter Notebooks, using Numpy 1.23^66^ and pandas 2.2^67^. Statistical tests were performed using the corresponding functions in Scipy 1.12^68^. Sequence data was processed with the help of Biopython 1.79^69^.

For all boxplots, boxes indicate the 25-75 percentile, whiskers extend to 1.5x the interquartile range, and outliers beyond this range are displayed as points.

## Acknowledgements

We would like to thank members of the Toth-Petroczy lab for useful comments and discussion. We would like to thank Andrej Shevchenko for advice on mass spectrometry-based proteomics. We thank Ksenia Kusnetsova and Swantje Lenz for critical reading of the manuscript.

We thank the Computer Department and the Scientific Computing Facility of the MPI of Molecular Cell Biology and Genetics for supporting our HPC system.

## References

1. Romero Romero, M. L., Landerer, C., Poehls, J. & Toth‐Petroczy, A. Phenotypic mutations contribute to protein diversity and shape protein evolution. Protein Science vol. 31 Preprint at 10.1002/pro.4397 (2022).

2. Goldsmith, M. & Tawfik, D. S. Potential role of phenotypic mutations in the evolution of protein expression and stability. Proc. Natl. Acad. Sci. U. S. A. 106, 6197–6202 (2009).

3. Bratulic, S., Toll-Riera, M. & Wagner, A. Mistranslation can enhance fitness through purging of deleterious mutations. Nat. Commun. 8, 15410 (2017).

4. Grosjean, H. J., de Henau, S. & Crothers, D. M. On the physical basis for ambiguity in genetic coding interactions. Proc. Natl. Acad. Sci. U. S. A. 75, 610–614 (1978).

5. Sharma, V. et al. Analysis of tetra- and hepta-nucleotides motifs promoting -1 ribosomal frameshifting in Escherichia coli. Nucleic Acids Res. 42, 7210–7225 (2014).

6. Romero Romero, M. L. et al. Environment modulates protein heterogeneity through transcriptional and translational stop codon readthrough. Nat. Commun. 15, 4446 (2024).

7. Čapková Pavlíková, Z. et al. Ribosomal A-site interactions with near-cognate tRNAs drive stop codon readthrough. Nat. Struct. Mol. Biol. (2025) doi:10.1038/s41594-024-01450-z.

8. Meyerovich, M., Mamou, G. & Ben-Yehuda, S. Visualizing high error levels during gene expression in living bacterial cells. Proc. Natl. Acad. Sci. U. S. A. 107, 11543–11548 (2010).

9. Bock, L. V. et al. Thermodynamic control of −1 programmed ribosomal frameshifting. Nat. Commun. 10, 1–11 (2019).

10. Kramer, E. B. & Farabaugh, P. J. The frequency of translational misreading errors in E. coli is largely determined by tRNA competition. RNA 13, 87–96 (2007).

11. Caliskan, N. et al. Conditional Switch between Frameshifting Regimes upon Translation of dnaX mRNA. Mol. Cell 66, 558-567.e4 (2017).

12. Curran, J. F. & Yarus, M. Use of tRNA suppressors to probe regulation of Escherichia coli release factor 2. J. Mol. Biol. 203, 75–83 (1988).

13. Pataskar, A. et al. Tryptophan depletion results in tryptophan-to-phenylalanine substitutants. Nature 603, 721–727 (2022).

14. Björk, G. R., Wikström, P. M. & Byström, A. S. Prevention of translational frameshifting by the modified nucleoside 1-methylguanosine. Science 244, 986–989 (1989).

15. Zaher, H. S. & Green, R. A primary role for release factor 3 in quality control during translation elongation in Escherichia coli. Cell 147, 396–408 (2011).

16. Peng, B.-Z. et al. Active role of elongation factor G in maintaining the mRNA reading frame during translation. Sci Adv 5, eaax8030 (2019).

17. Jacks, T. et al. Characterization of ribosomal frameshifting in HIV-1 gag-pol expression. Nature 331, 280–283 (1988).

18. Yanagida, H. et al. The Evolutionary Potential of Phenotypic Mutations. PLoS Genet. 11, e1005445 (2015).

19. Meydan, S. et al. Programmed Ribosomal Frameshifting Generates a Copper Transporter and a Copper Chaperone from the Same Gene. Mol. Cell 65, 207–219 (2017).

20. Craigen, W. J. & Caskey, C. T. Expression of peptide chain release factor 2 requires high-efficiency frameshift. Nature 322, 273–275 (1986).

21. Plant, E. P., Rakauskaite, R., Taylor, D. R. & Dinman, J. D. Achieving a golden mean: mechanisms by which coronaviruses ensure synthesis of the correct stoichiometric ratios of viral proteins. J. Virol. 84, 4330–4340 (2010).

22. Tsuchihashi, Z. & Kornberg, A. Translational frameshifting generates the gamma subunit of DNA polymerase III holoenzyme. Proc. Natl. Acad. Sci. U. S. A. (1990).

23. Ivanov, I. P., Simin, K., Letsou, A., Atkins, J. F. & Gesteland, R. F. The Drosophila gene for antizyme requires ribosomal frameshifting for expression and contains an intronic gene for snRNP Sm D3 on the opposite strand. Mol. Cell. Biol. 18, 1553–1561 (1998).

24. Jungreis, I. et al. Evidence of abundant stop codon readthrough in Drosophila and other metazoa. Genome Res. 21, 2096–2113 (2011).

25. Stiebler, A. C. et al. Ribosomal readthrough at a short UGA stop codon context triggers dual localization of metabolic enzymes in Fungi and animals. PLoS Genet. 10, e1004685 (2014).

26. Kachale, A. et al. Short tRNA anticodon stem and mutant eRF1 allow stop codon reassignment. Nature 613, 751–758 (2023).

27. Namy, O., Duchateau-Nguyen, G. & Rousset, J.-P. Translational readthrough of the PDE2 stop codon modulates cAMP levels in Saccharomyces cerevisiae. Mol. Microbiol. 43, 641–652 (2002).

28. Schueren, F. et al. Peroxisomal lactate dehydrogenase is generated by translational readthrough in mammals. Elife 3, (2014).

29. Landerer, C., Pöhls, J. & Toth-Petroczy, A. Fitness effects of phenotypic mutations at proteome-scale reveal optimality of translation machinery. Mol. Biol. Evol. 41, (2024).

30. Miranda, I. et al. Candida albicans CUG mistranslation is a mechanism to create cell surface variation. MBio 4, (2013).

31. Netzer, N. et al. Innate immune and chemically triggered oxidative stress modifies translational fidelity. Nature 462, 522–526 (2009).

32. Savitski, M. M., Nielsen, M. L. & Zubarev, R. A. ModifiComb, a new proteomic tool for mapping substoichiometric post-translational modifications, finding novel types of modifications, and fingerprinting complex protein mixtures. Mol. Cell. Proteomics 5, 935–948 (2006).

33. Yu, F. et al. Identification of modified peptides using localization-aware open search. Nat. Commun. 11, 4065 (2020).

34. Cox, J. & Mann, M. MaxQuant enables high peptide identification rates, individualized p.p.b.-range mass accuracies and proteome-wide protein quantification. Nat. Biotechnol. 26, 1367–1372 (2008).

35. Mordret, E. et al. Systematic detection of amino acid substitutions in proteomes reveals mechanistic basis of ribosome errors and selection for translation fidelity. Mol. Cell 75, 427-441.e5 (2019).

36. Wu, X., Xu, M., Yang, J.-R. & Lu, J. Genome-wide impact of codon usage bias on translation optimization in Drosophila melanogaster. Nat. Commun. 15, 8329 (2024).

37. Perez-Riverol, Y. et al. The PRIDE database at 20 years: 2025 update. Nucleic Acids Res. 53, D543–D553 (2025).

38. Kong, A. T., Leprevost, F. V., Avtonomov, D. M., Mellacheruvu, D. & Nesvizhskii, A. I. MSFragger: Ultrafast and comprehensive peptide identification in mass spectrometry-based proteomics. Nat. Methods 14, 513–520 (2017).

39. Sun, L. et al. Evolutionary gain of alanine mischarging to noncognate tRNAs with a G4:U69 base pair. J. Am. Chem. Soc. 138, 12948–12955 (2016).

40. Rozov, A. et al. Novel base-pairing interactions at the tRNA wobble position crucial for accurate reading of the genetic code. Nat. Commun. 7, 1–10 (2016).

41. Joshi, K., Bhatt, M. J. & Farabaugh, P. J. Codon-specific effects of tRNA anticodon loop modifications on translational misreading errors in the yeast Saccharomyces cerevisiae. Nucleic Acids Res. 46, 10331–10339 (2018).

42. Rozov, A., Demeshkina, N., Westhof, E., Yusupov, M. & Yusupova, G. Structural insights into the translational infidelity mechanism. Nat. Commun. 6, 7251 (2015).

43. Edelmann, P. & Gallant, J. Mistranslation in E. coli. Cell 10, 131–137 (1977).

44. Parker, J. & Friesen, J. D. “Two out of three” codon reading leading to mistranslation in vivo. Mol. Gen. Genet. 177, 439–445 (1980).

45. Bouadloun, F., Donner, D. & Kurland, C. G. Codon-specific missense errors in vivo. EMBO J. 2, 1351–1356 (1983).

46. Stikeleather, R., Ali, F., Ho, W.-C., Licknack, T. & Lynch, M. Translation accuracy in E. coli. bioRxivorg (2025) doi:10.1101/2025.04.18.649569.

47. Tretyachenko, V. et al. Encoded and non-genetic alternative protein variants expand human functional proteome. bioRxiv (2025) doi:10.1101/2025.02.17.638604.

48. Tsour, S. et al. Alternate RNA decoding results in stable and abundant proteins in mammals. bioRxivorg (2024) doi:10.1101/2024.08.26.609665.

49. Michalski, A., Cox, J. & Mann, M. More than 100,000 detectable peptide species elute in single shotgun proteomics runs but the majority is inaccessible to data-dependent LC-MS/MS. J. Proteome Res. 10, 1785–1793 (2011).

50. Cech, N. B. & Enke, C. G. Relating electrospray ionization response to nonpolar character of small peptides. Anal. Chem. 72, 2717–2723 (2000).

51. Liigand, P., Kaupmees, K. & Kruve, A. Influence of the amino acid composition on the ionization efficiencies of small peptides. J. Mass Spectrom. 54, 481–487 (2019).

52. Yang, C. et al. Arginine deprivation enriches lung cancer proteomes with cysteine by inducing arginine-to-cysteine substitutants. Mol. Cell 84, 1904-1916.e7 (2024).

53. Kubo, T. et al. Active site cysteine-null glyceraldehyde-3-phosphate dehydrogenase (GAPDH) rescues nitric oxide-induced cell death. Nitric Oxide 53, 13–21 (2016).

54. Nakajima, H. et al. The active site cysteine of the proapoptotic protein glyceraldehyde-3-phosphate dehydrogenase is essential in oxidative stress-induced aggregation and cell death. J. Biol. Chem. 282, 26562–26574 (2007).

55. Whitehead, D. J., Wilke, C. O., Vernazobres, D. & Bornberg-Bauer, E. The look-ahead effect of phenotypic mutations. Biol. Direct 3, 18 (2008).

56. Kurland, C. G. & Ehrenberg, M. Optimization of translation accuracy. Prog. Nucleic Acid Res. Mol. Biol. 31, 191–219 (1984).

57. Wohlgemuth, I., Pohl, C., Mittelstaet, J., Konevega, A. L. & Rodnina, M. V. Evolutionary optimization of speed and accuracy of decoding on the ribosome. Philos. Trans. R. Soc. Lond. B Biol. Sci. 366, 2979–2986 (2011).

58. Hopfield, J. J. Kinetic proofreading: a new mechanism for reducing errors in biosynthetic processes requiring high specificity. Proc. Natl. Acad. Sci. U. S. A. 71, 4135–4139 (1974).

59. Ruusala, T., Andersson, D., Ehrenberg, M. & Kurland, C. G. Hyper-accurate ribosomes inhibit growth. EMBO J. 3, 2575–2580 (1984).

60. Chang, H.-Y. et al. Crystal-C: A Computational Tool for Refinement of Open Search Results. J. Proteome Res. 19, 2511–2515 (2020).

61. Keller, A., Nesvizhskii, A. I., Kolker, E. & Aebersold, R. Empirical statistical model to estimate the accuracy of peptide identifications made by MS/MS and database search. Anal. Chem. 74, 5383–5392 (2002).

62. Nesvizhskii, A. I., Keller, A., Kolker, E. & Aebersold, R. A statistical model for identifying proteins by tandem mass spectrometry. Anal. Chem. 75, 4646–4658 (2003).

63. da Veiga Leprevost, F. et al. Philosopher: a versatile toolkit for shotgun proteomics data analysis. Nat. Methods 17, 869–870 (2020).

64. Benjamini, Y. & Hochberg, Y. Controlling the false discovery rate: A practical and powerful approach to multiple testing. J. R. Stat. Soc. 57, 289–300 (1995).

65. VanRossum, G. & Drake, F. L. The python language reference. https://scicomp.ethz.ch/public/manual/Python/3.9./reference.pdf.

66. Harris, C. R. et al. Array programming with NumPy. Nature 585, 357–362 (2020).

67. Pandas Team. Pandas-Dev/Pandas: Pandas. (Zenodo, 2024). doi:10.5281/zenodo.10697587.

68. Virtanen, P. et al. SciPy 1.0: fundamental algorithms for scientific computing in Python. Nat. Methods 17, 261–272 (2020).

69. Cock, P. J. A. et al. Biopython: freely available Python tools for computational molecular biology and bioinformatics. Bioinformatics 25, 1422–1423 (2009).

