## Supplementary Information for "Amino acid and codon usage explain amino acid misincorporation rates across the tree of life"

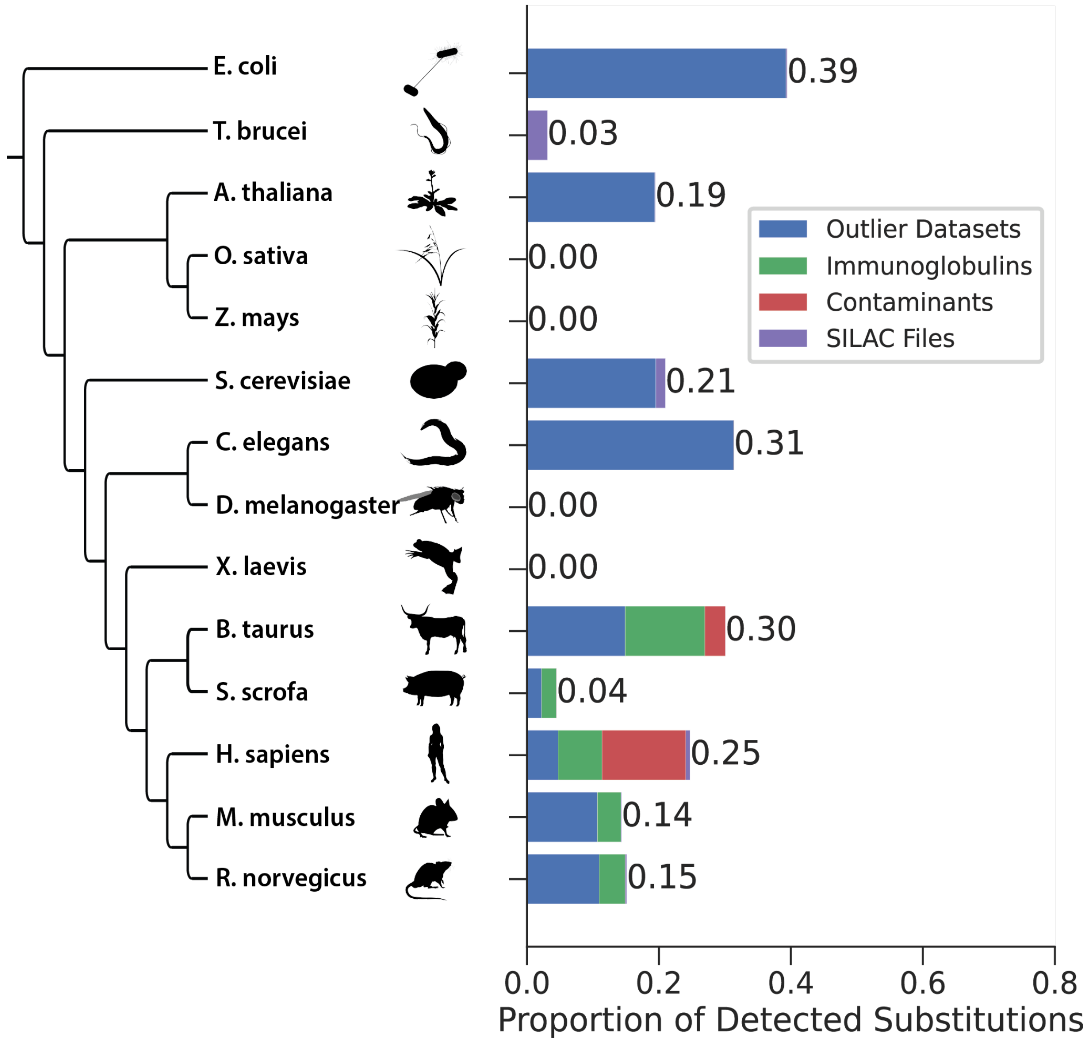


**Supplementary Figure 1: Filtering of the datasets by removing outliers and contaminants.** Proportions of originally detected misincorporations that were removed from all analyses because the dataset had unusually high error rates, the misincorporation mapped to an immunoglobulin, to a common proteomics contaminant protein, or the MS raw file showed a high proportion of SILAC-like mass shifts.

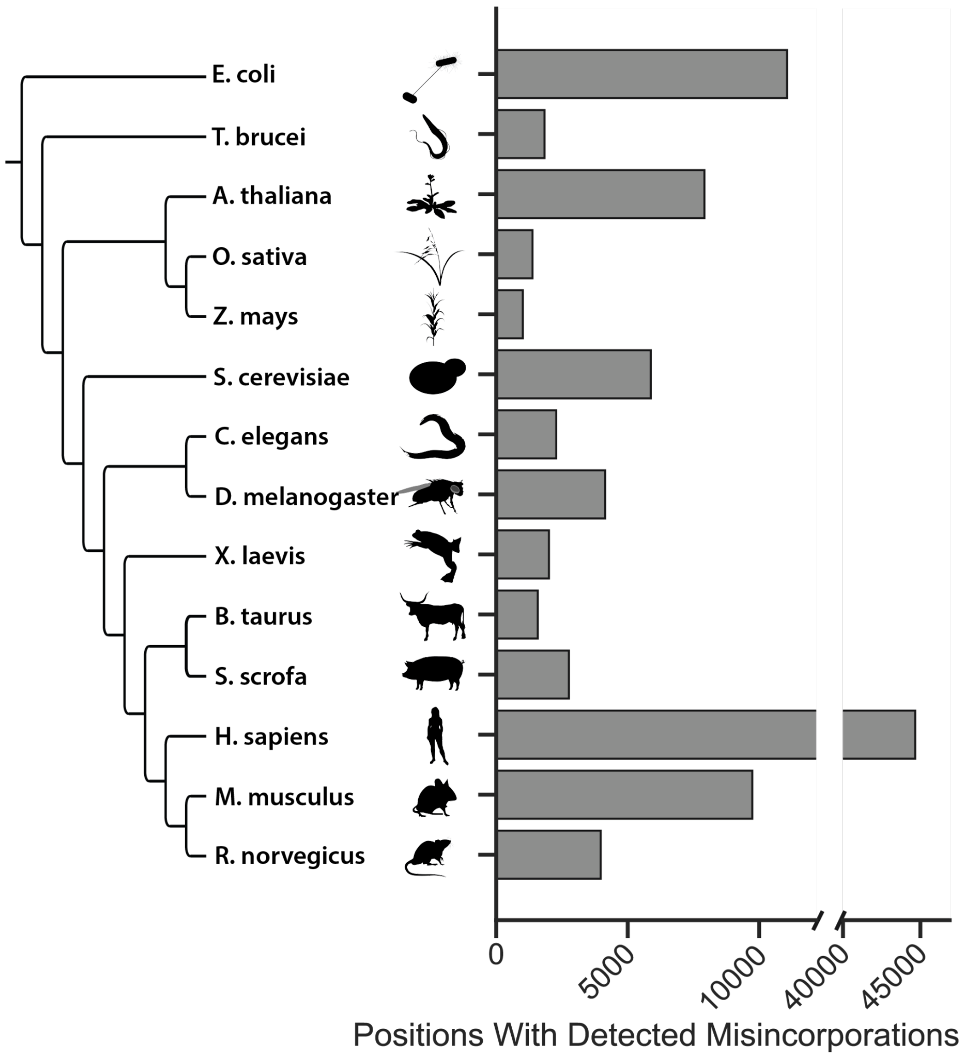


**Supplementary Figure 2: Number of different proteome positions with at least one detected misincorporation, per species.**

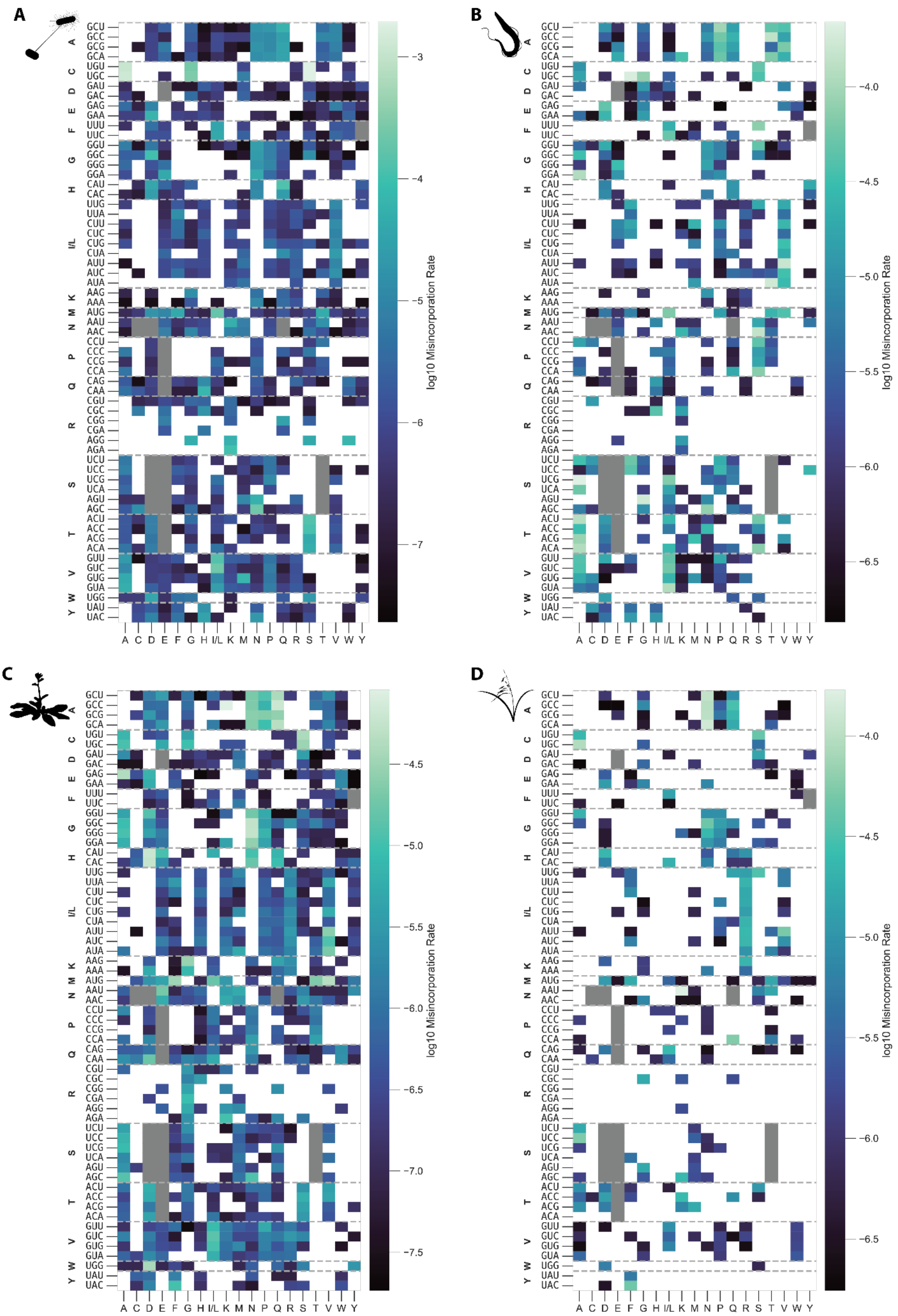
A
B
C
D
GCU GCC GCG GCA UGU UGC GAU GAC GAG GAA UUU UUC GGU GGC GGG GGA CAU CAC UUG UUA CUU CUC CUG CUA AUU AUC AUA AAG AAA AUG AAU AAC CCU CCC CCG CCA CAG CAA CGU CGC CGG CGA AGG AGA UCU UCC UCG UCA AGU AGC ACU ACC ACG ACA GUU GUC GUG GUA UGG UAU UAC
A C D E F G H I/L K M N P Q R S T V W Y
log10 Misincorporation Rate
−3
−4
−5
−6
−7
−4.0
−4.5
−5.0
−5.5
−6.0
−6.5
−4.5
−5.0
−5.5
−6.0
−6.5
−7.0
−7.5
−4.0
−4.5
−5.0
−5.5
−6.0
−6.5

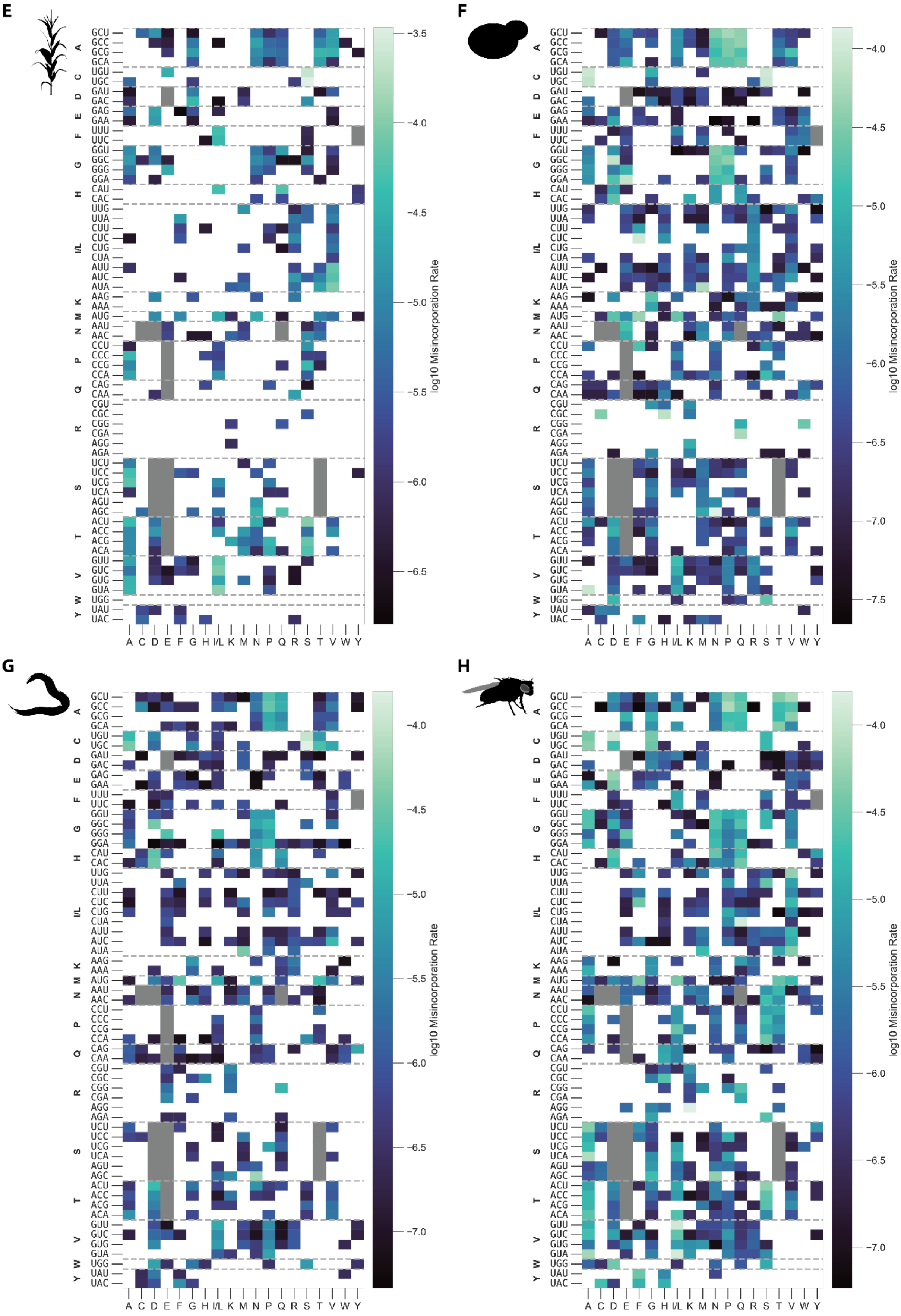
E
F
G
H
log10 Misincorporation Rate
A C D E F G H I/L K M N P Q R S T V W Y
A C D E F G H I/L K M N P Q R S T V W Y
A C D E F G H I/L K M N P Q R S T V W Y
A C D E F G H I/L K M N P Q R S T V W Y

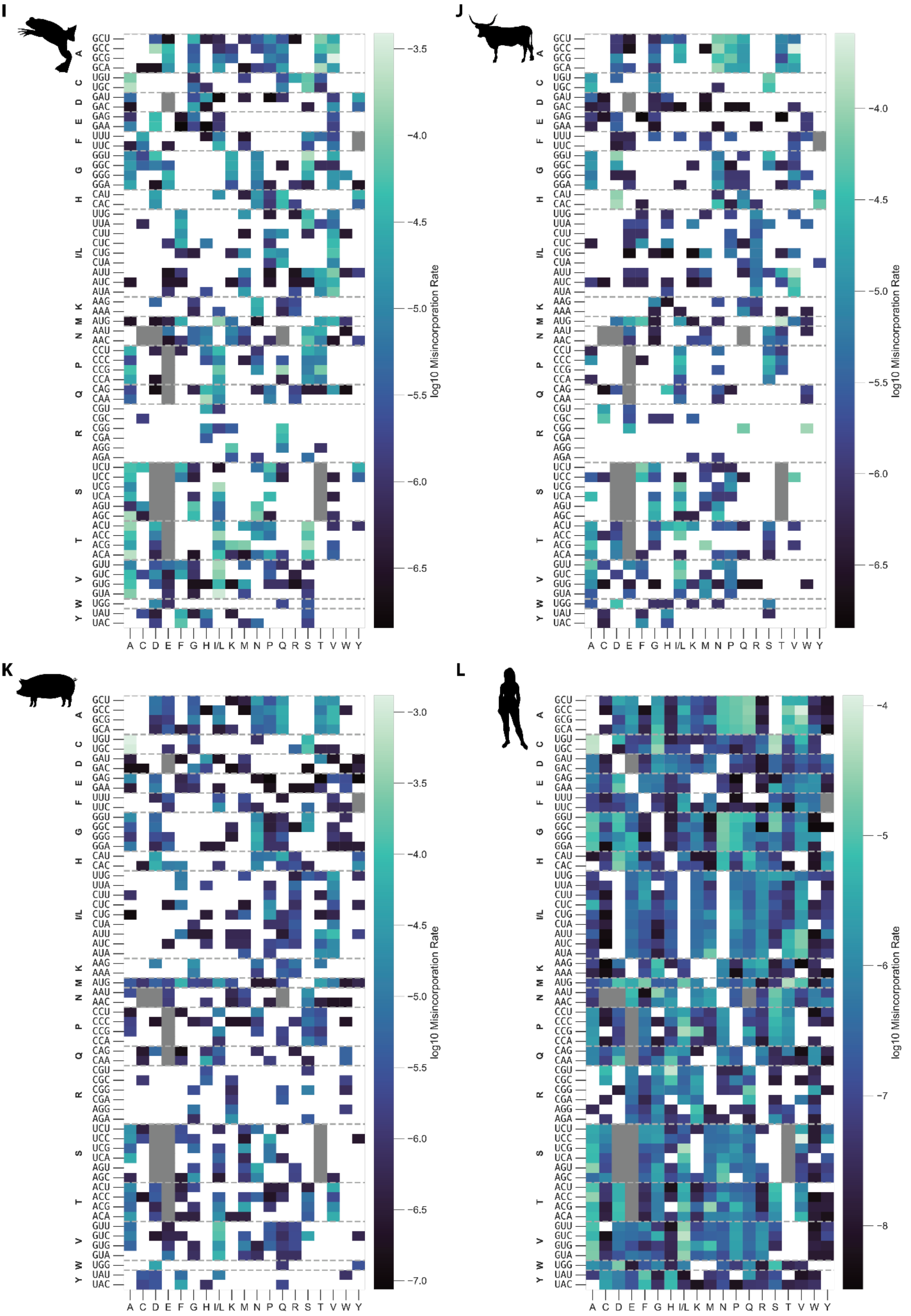
I
J
K
L
GCU GCC GCG GCA UGU UGC GAU GAC GAG GAA UUU UUC GGU GGC GGG GGA CAU CAC UUG UUA CUU CUC CUG CUA AUU AUC AUA AAG AAA AUG AAU AAC CCU CCC CCG CCA CAG CAA CGU CGC CGG CGA AGG AGA UCU UCC UCG UCA AGU AGC ACU ACC ACG ACA GUU GUC GUG GUA UGG UAU UAC
A C D E F G H I/L K M N P Q R S T V W Y
log10 Misincorporation Rate
−3.5 −4.0 −4.5 −5.0 −5.5 −6.0 −6.5
−4.0 −4.5 −5.0 −5.5 −6.0 −6.5
−3.0 −3.5 −4.0 −4.5 −5.0 −5.5 −6.0 −6.5 −7.0
−4 −5 −6 −7 −8

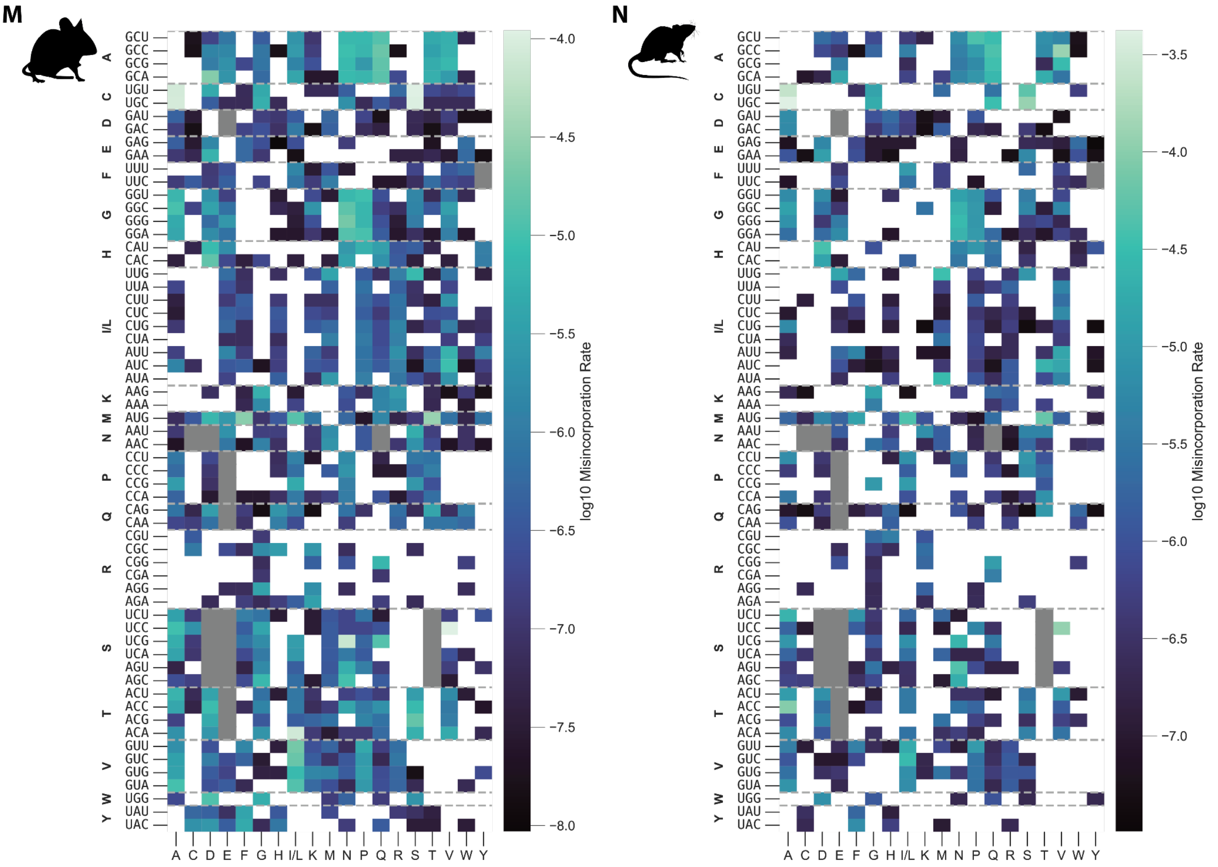


**Supplementary Figure 3: Codon-to-amino acid misincorporation rates for each species.** Shown are the logarithmic misincorporation rates from each codon to each amino acid by species. Each field corresponds to the number of times the given codon was observed (PSMs) with a misincorporation of the given amino acid, divided by the total number of observations (spectral counts) of that codon. White fields correspond to misincorporations not observed in the given species. Gray fields indicate misincorporations undetectable due to a mass shift that is equal or very similar to a PTM. A: *E. coli*, B: *T. brucei*, C: *A. thaliana*, D: *O. sativa*, E: *Z. mays*, F: *S. cerevisiae*, G: *C. elegans*, H: *D. melanogaster*, I: *X. laevis*, J: *B. taurus*, K: *S. scrofa*, L: *H. sapiens*, M: *M. musculus*, N: *R. norvegicus*.

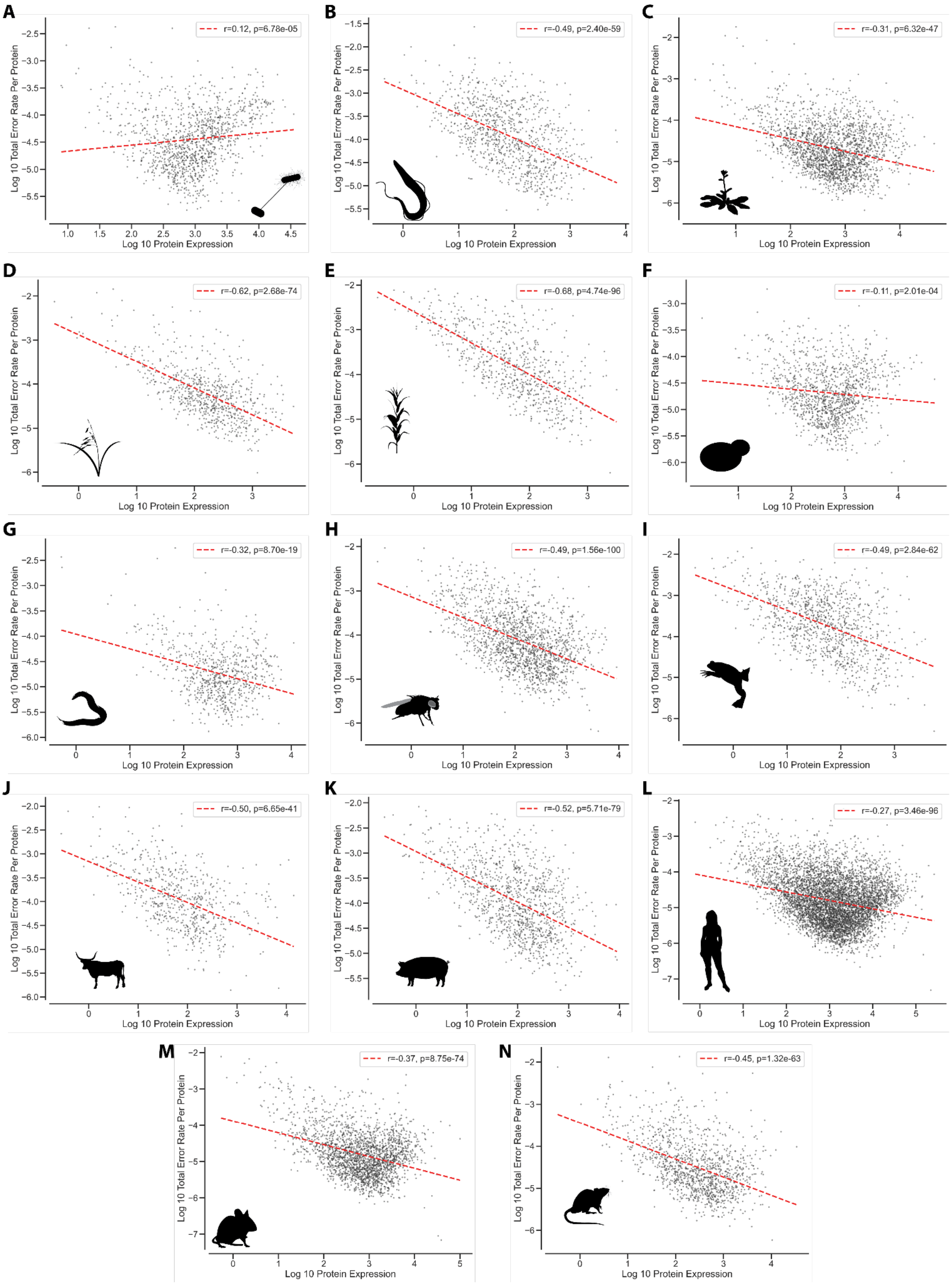


**Supplementary Figure 4: Correlation of per-protein misincorporation rates and protein expression level**. Protein expression is equal to the total spectral count of all peptides mapping to the protein, divided by the protein's length. r: Pearson correlation coefficient. P-values calculated using two-sided Wald test. A: *E. coli*, B: *T. brucei*, C: *A. thaliana*, D: *O. sativa*,

E: *Z. mays*, F: *S. cerevisiae*, G: *C. elegans*, H: *D. melanogaster*, I: *X. laevis*, J: *B. taurus*, K: *S. scrofa*, L: *H. sapiens*, M: *M. musculus*, N: *R. norvegicus*.

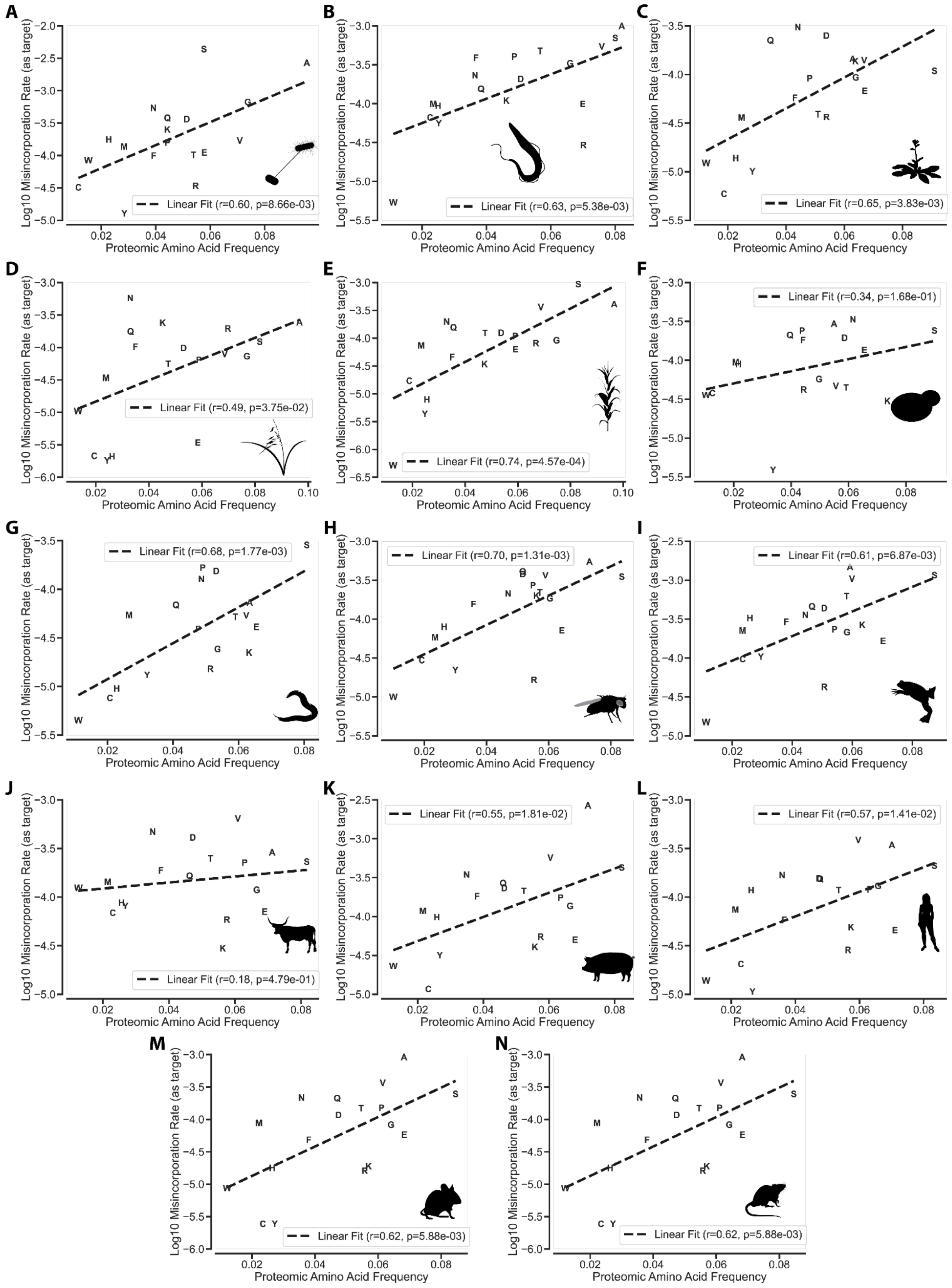

**Supplementary Figure 5: Correlation of frequency of amino acids and rate of misincorporation.** Amino acid frequency refers to the proportion of positions in the proteome occupied by that amino acid. r: Pearson correlation coefficient. P-values calculated using two-sided Wald test. A: *E. coli*, B: *T. brucei*, C: *A. thaliana*, D: *O. sativa*, E: *Z. mays*, F: *S. cerevisiae*, G: *C. elegans*, H: *D. melanogaster*, I: *X. laevis*, J: *B. taurus*, K: *S. scrofa*, L: *H. sapiens*, M: *M. musculus*, N: *R. norvegicus*.

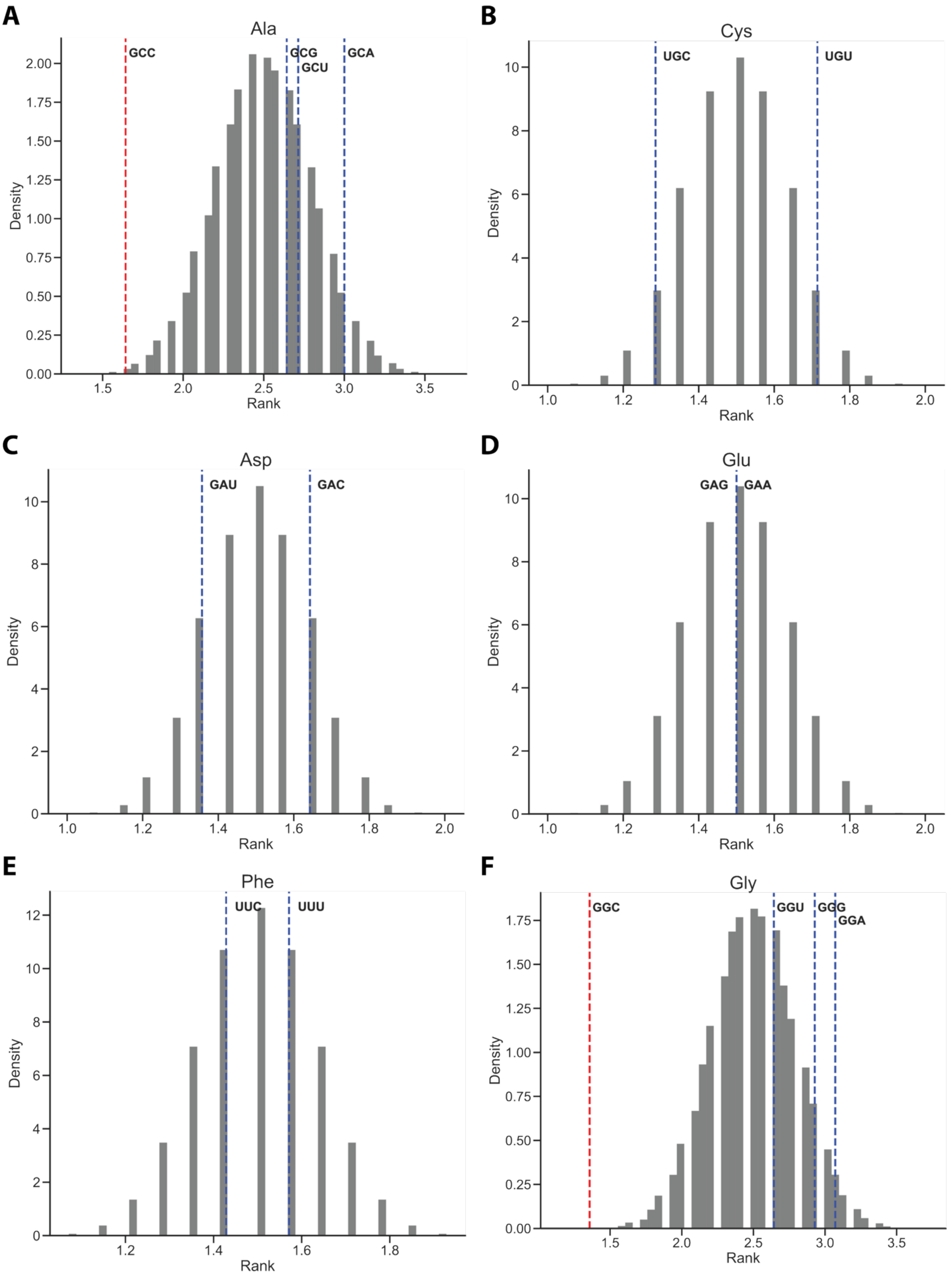
A
Ala
GCC
GCG
GCU
GCA
Density
Rank
B
Cys
UGC
UGU
Density
Rank
C
Asp
GAU
GAC
Density
Rank
D
Glu
GAG
GAA
Density
Rank
E
Phe
UUC
UUU
Density
Rank
F
Gly
GGC
GGU
GGG
GGA
Density
Rank

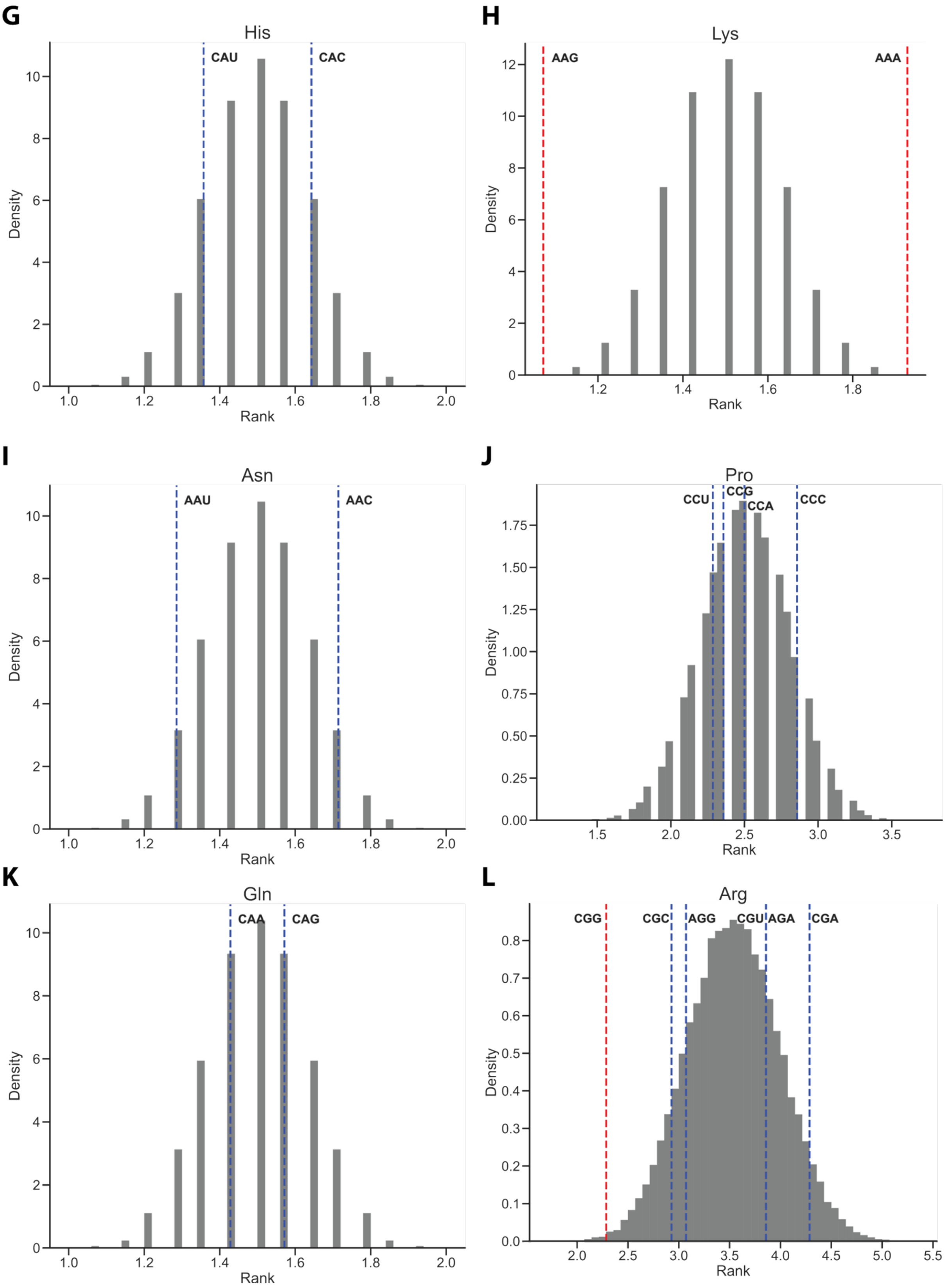
G
His
CAU
CAC
Density
Rank
H
Lys
AAG
AAA
Density
Rank
I
Asn
AAU
AAC
Density
Rank
J
Pro
CCU
CCG
CCA
CCC
Density
Rank
K
Gln
CAA
CAG
Density
Rank
L
Arg
CGG
CGC
AGG
CGU
AGA
CGA
Density
Rank

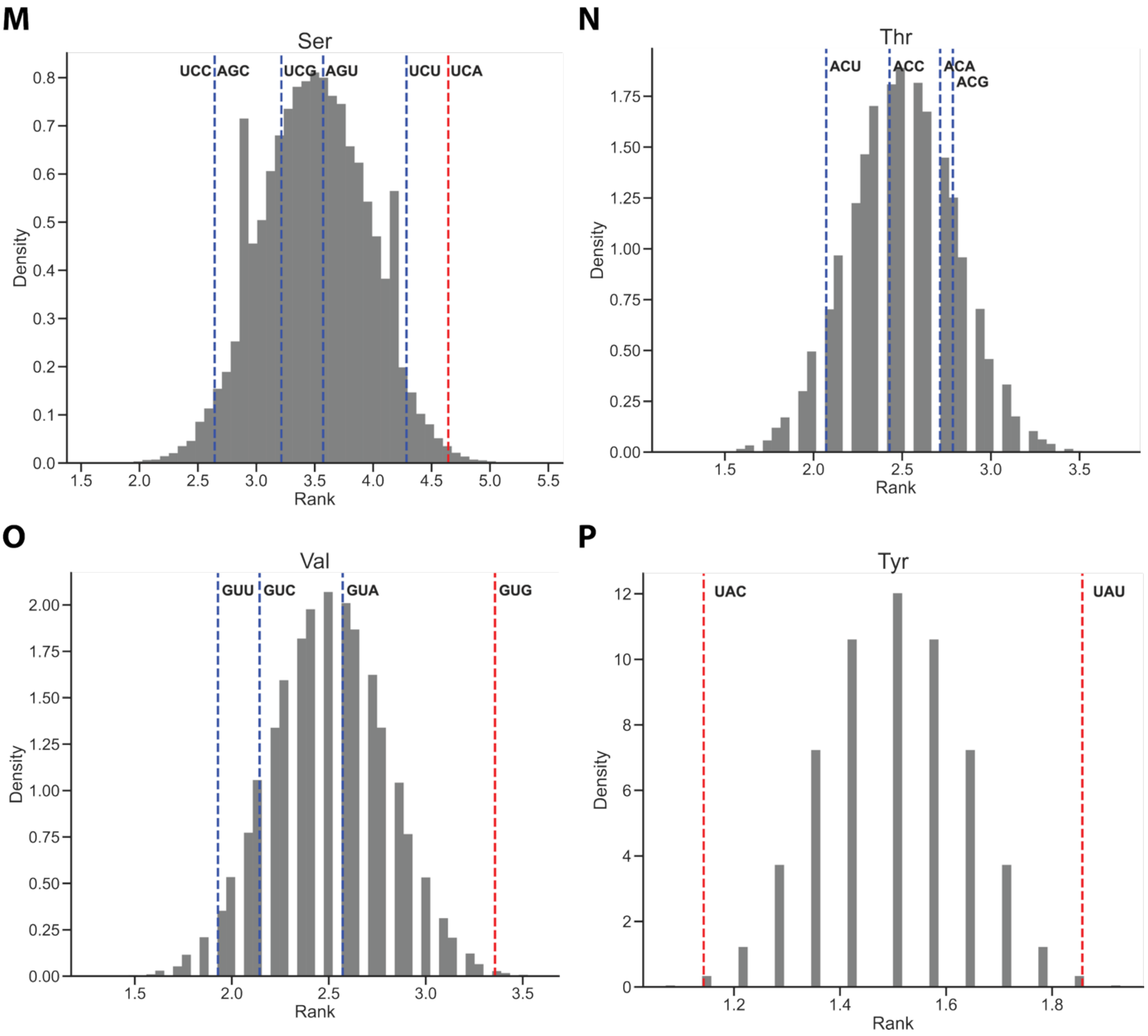


**Supplementary Figure 6: Mean error rate ranks of synonymous codons vs bootstrapped expectation.** Within each species, for each amino acid, the synonymous codons were ranked according to their total error rate (rank 1 means highest error rate). For each amino acid, 10,000 sets of ranks were randomly generated (gray histogram). The mean rank across species of each codon was then compared to this random distribution, and an empirical p-value calculated. Red lines indicate p<0.05 (one-sided). For Fig 3C, p-values were adjusted for multiple testing by Benjamini-Hochberg method.

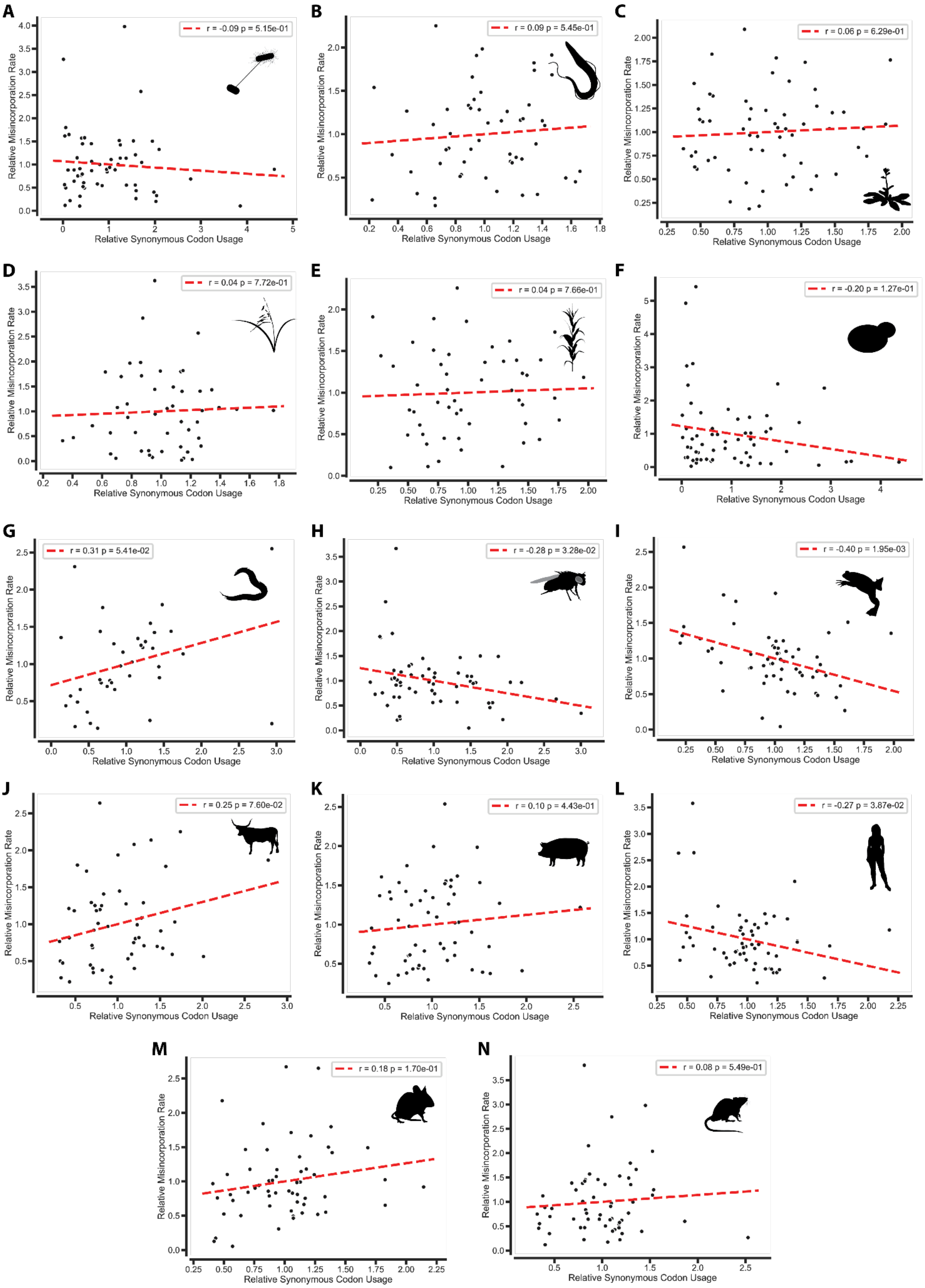


**Supplementary Figure 7: Correlation between relative synonymous codon usage (RSCU) and relative misincorporation rate (RMR) for all species.** RMR is a codon's total error rate divided by the mean total error rate of its synonyms. r: Pearson correlation coefficient. P-values calculated using two-sided Wald test. A: *E. coli*, B: *T. brucei*, C: *A.*

*thaliana*, D: *O. sativa*, E: *Z. mays*, F: *S. cerevisiae*, G: *C. elegans*, H: *D. melanogaster*, I: *X. laevis*, J: *B. taurus*, K: *S. scrofa*, L: *H. sapiens*, M: *M. musculus*, N: *R. norvegicus*.

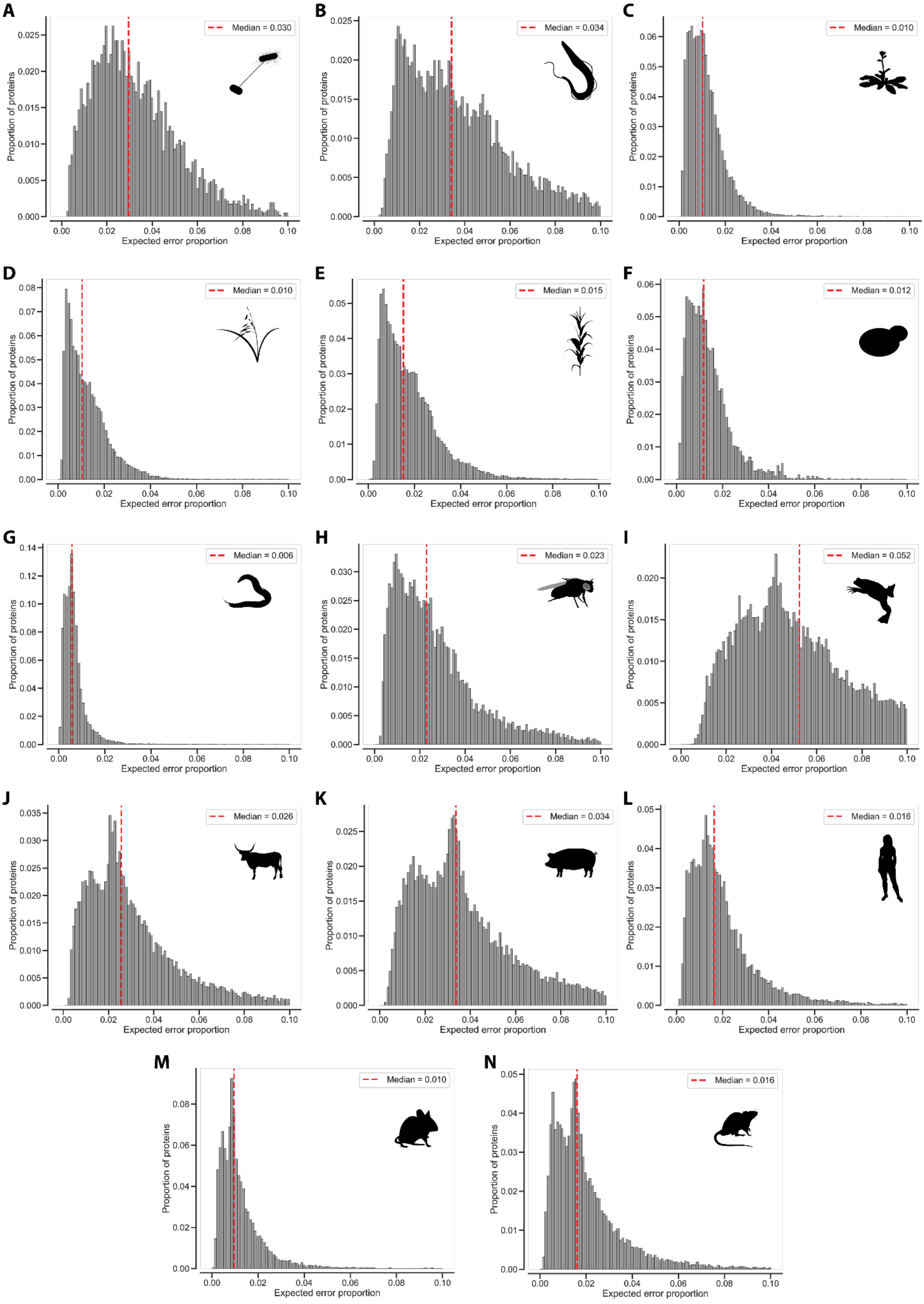

**Supplementary Figure 8: Expected proportion of protein molecules containing at least one misincorporation.** Histogram of error proportion for all proteins in the respective proteome. Red dotted line indicates median value. A: *E. coli*, B: *T. brucei*, C: *A. thaliana*, D: *O. sativa*, E: *Z. mays*, F: *S. cerevisiae*, G: *C. elegans*, H: *D. melanogaster*, I: *X. laevis*, J: *B. taurus*, K: *S. scrofa*, L: *H. sapiens*, M: *M. musculus*, N: *R. norvegicus*.

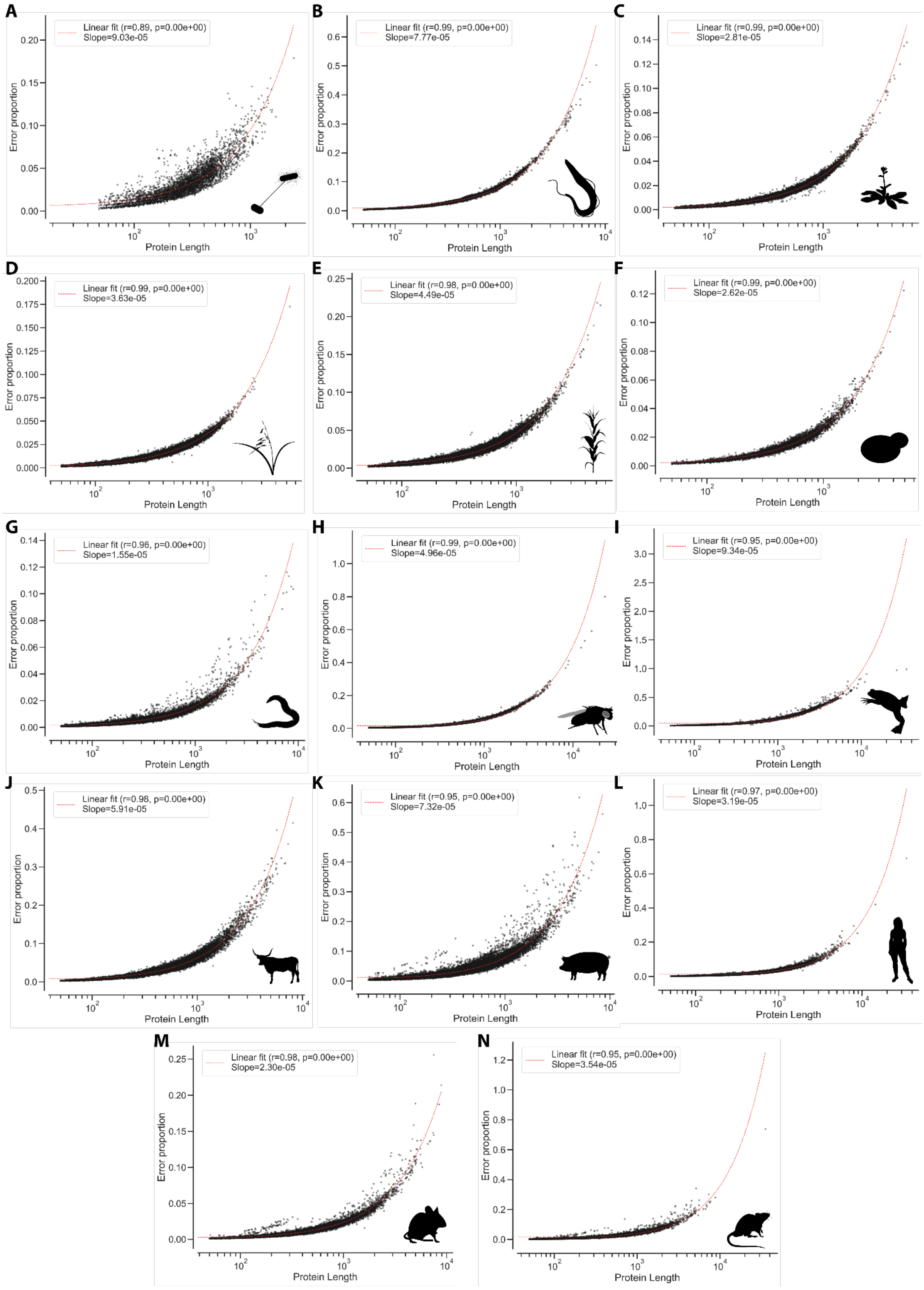


**Supplementary Figure 9: Correlation between protein length and error proportion for all species.** r: Pearson correlation coefficient. P-values calculated using two-sided Wald test. A: E. coli, B: T. brucei, C: A. thaliana, D: O. sativa, E: Z. mays, F: S. cerevisiae, G: C. elegans,

H: D. melanogaster, I: X. laevis, J: B. taurus, K: S. scrofa, L: H. sapiens, M: M. musculus, N: R. norvegicus.

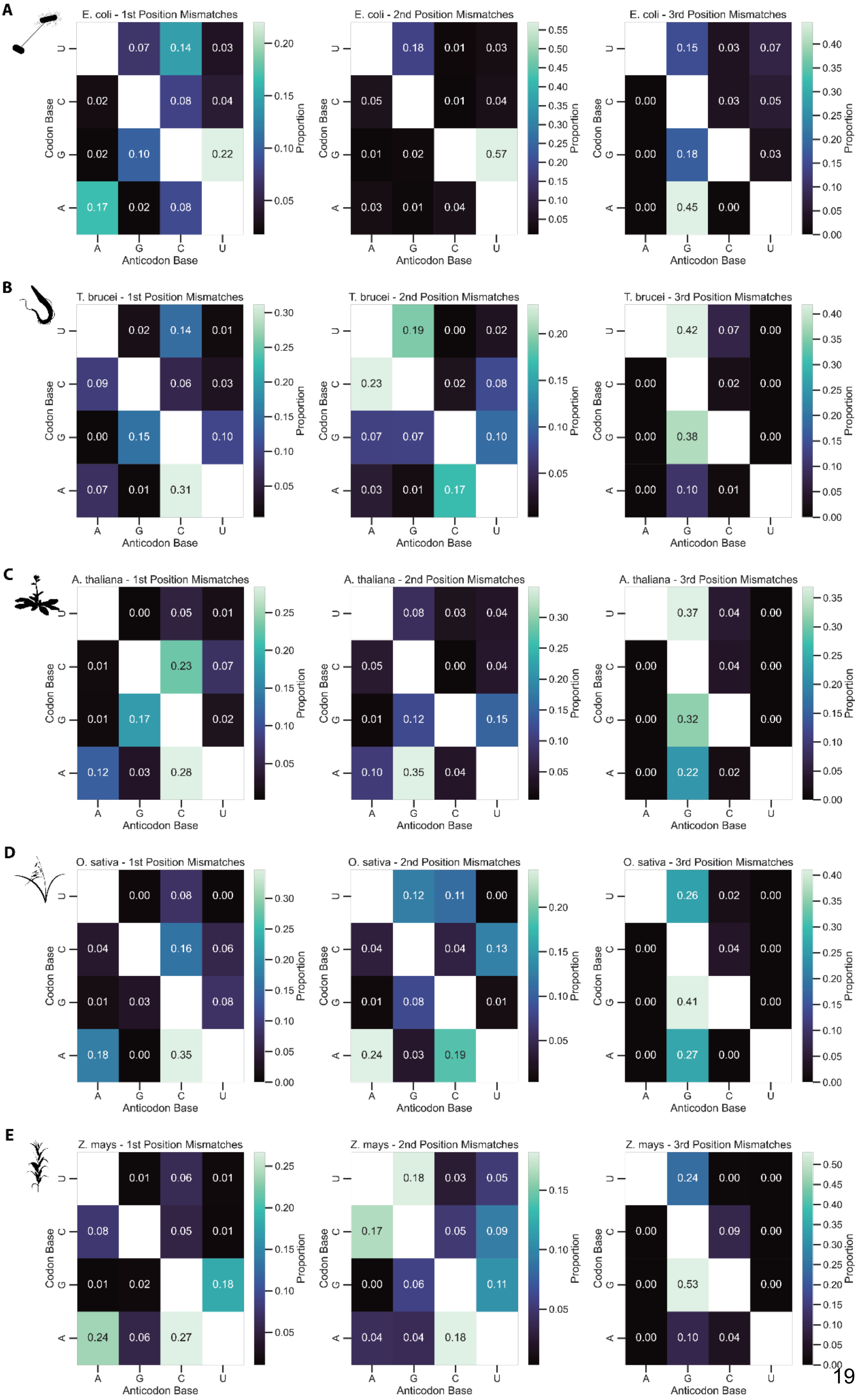

A
E. coli - 1st Position Mismatches
E. coli - 2nd Position Mismatches
E. coli - 3rd Position Mismatches
B
T. brucei - 1st Position Mismatches
T. brucei - 2nd Position Mismatches
T. brucei - 3rd Position Mismatches
C
A. thaliana - 1st Position Mismatches
A. thaliana - 2nd Position Mismatches
A. thaliana - 3rd Position Mismatches
D
O. sativa - 1st Position Mismatches
O. sativa - 2nd Position Mismatches
O. sativa - 3rd Position Mismatches
E
Z. mays - 1st Position Mismatches
Z. mays - 2nd Position Mismatches
Z. mays - 3rd Position Mismatches
Codon Base
Anticodon Base
Proportion

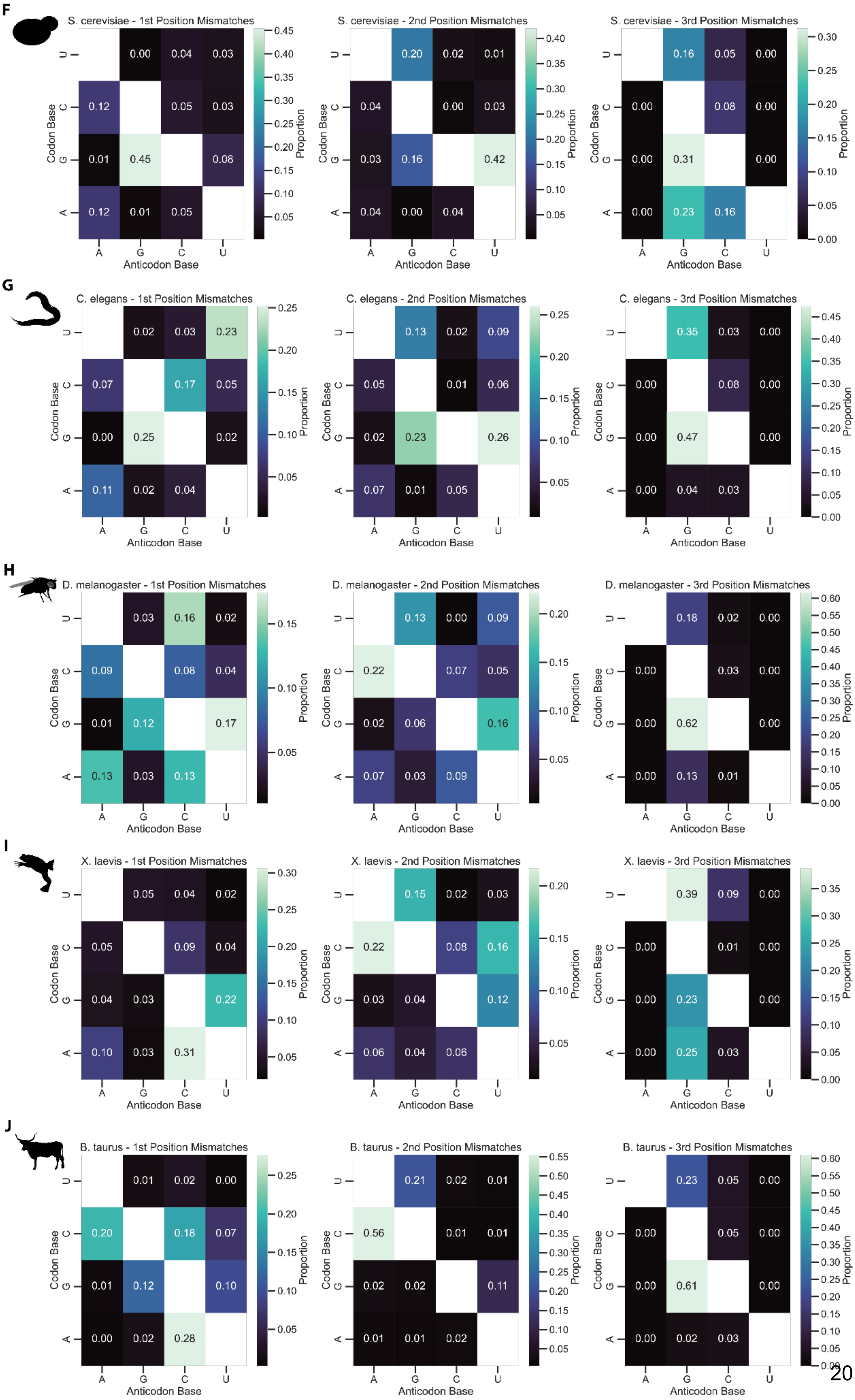
F
S. cerevisiae - 1st Position Mismatches
U 0.00 0.04 0.03
C 0.12 0.05 0.03
G 0.01 0.45 0.08
A 0.12 0.01 0.05
S. cerevisiae - 2nd Position Mismatches
U 0.20 0.02 0.01
C 0.04 0.00 0.03
G 0.03 0.16 0.42
A 0.04 0.00 0.04
S. cerevisiae - 3rd Position Mismatches
U 0.16 0.05 0.00
C 0.00 0.08 0.00
G 0.00 0.31 0.00
A 0.00 0.23 0.16
G
C. elegans - 1st Position Mismatches
U 0.02 0.03 0.23
C 0.07 0.17 0.05
G 0.00 0.25 0.02
A 0.11 0.02 0.04
C. elegans - 2nd Position Mismatches
U 0.13 0.02 0.09
C 0.05 0.01 0.06
G 0.02 0.23 0.26
A 0.07 0.01 0.05
C. elegans - 3rd Position Mismatches
U 0.35 0.03 0.00
C 0.00 0.08 0.00
G 0.00 0.47 0.00
A 0.00 0.04 0.03
H
D. melanogaster - 1st Position Mismatches
U 0.03 0.16 0.02
C 0.09 0.08 0.04
G 0.01 0.12 0.17
A 0.13 0.03 0.13
D. melanogaster - 2nd Position Mismatches
U 0.13 0.00 0.09
C 0.22 0.07 0.05
G 0.02 0.06 0.16
A 0.07 0.03 0.09
D. melanogaster - 3rd Position Mismatches
U 0.18 0.02 0.00
C 0.00 0.03 0.00
G 0.00 0.62 0.00
A 0.00 0.13 0.01
I
X. laevis - 1st Position Mismatches
U 0.05 0.04 0.02
C 0.05 0.09 0.04
G 0.04 0.03 0.22
A 0.10 0.03 0.31
X. laevis - 2nd Position Mismatches
U 0.15 0.02 0.03
C 0.22 0.08 0.16
G 0.03 0.04 0.12
A 0.06 0.04 0.06
X. laevis - 3rd Position Mismatches
U 0.39 0.09 0.00
C 0.00 0.01 0.00
G 0.00 0.23 0.00
A 0.00 0.25 0.03
J
B. taurus - 1st Position Mismatches
U 0.01 0.02 0.00
C 0.20 0.18 0.07
G 0.01 0.12 0.10
A 0.00 0.02 0.28
B. taurus - 2nd Position Mismatches
U 0.21 0.02 0.01
C 0.56 0.01 0.01
G 0.02 0.02 0.11
A 0.01 0.01 0.02
B. taurus - 3rd Position Mismatches
U 0.23 0.05 0.00
C 0.00 0.05 0.00
G 0.00 0.61 0.00
A 0.00 0.02 0.03
Codon Base
Anticodon Base
A G C U
Proportion

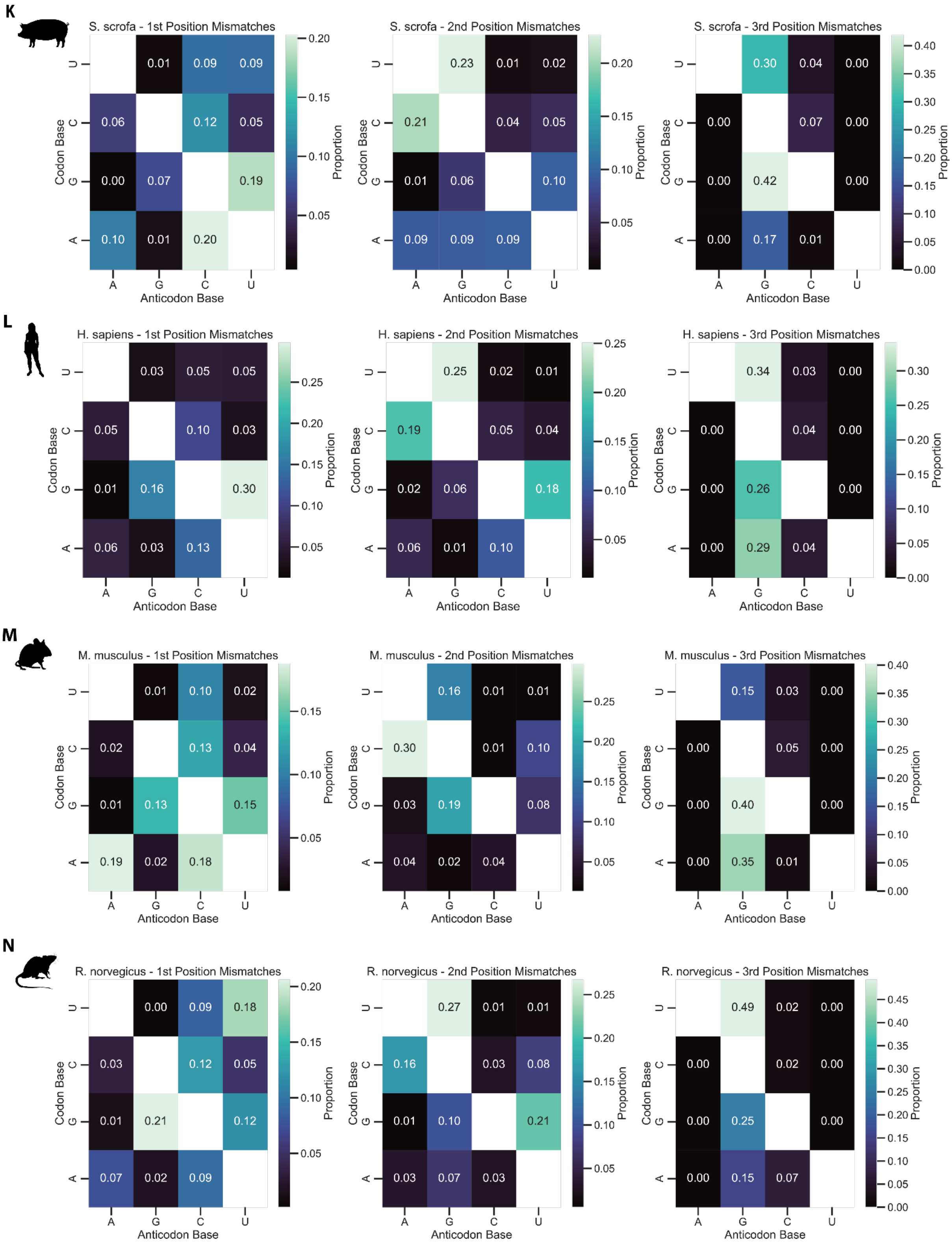


**Supplementary Figure 10: Proportions of inferred nucleotide mismatches for likely mispairing misincorporations by codon position, for each species.** Proportions are relative to all misincorporations classified as likely mispairing that have a mismatch at the respective codon position. A: *E. coli*, B: *T. brucei*, C: *A. thaliana*, D: *O. sativa*, E: *Z. mays*, F: *S. cerevisiae*, G: *C. elegans*, H: *D. melanogaster*, I: *X. laevis*, J: *B. taurus*, K: *S. scrofa*, L: *H. sapiens*, M: *M. musculus*, N: *R. norvegicus*.

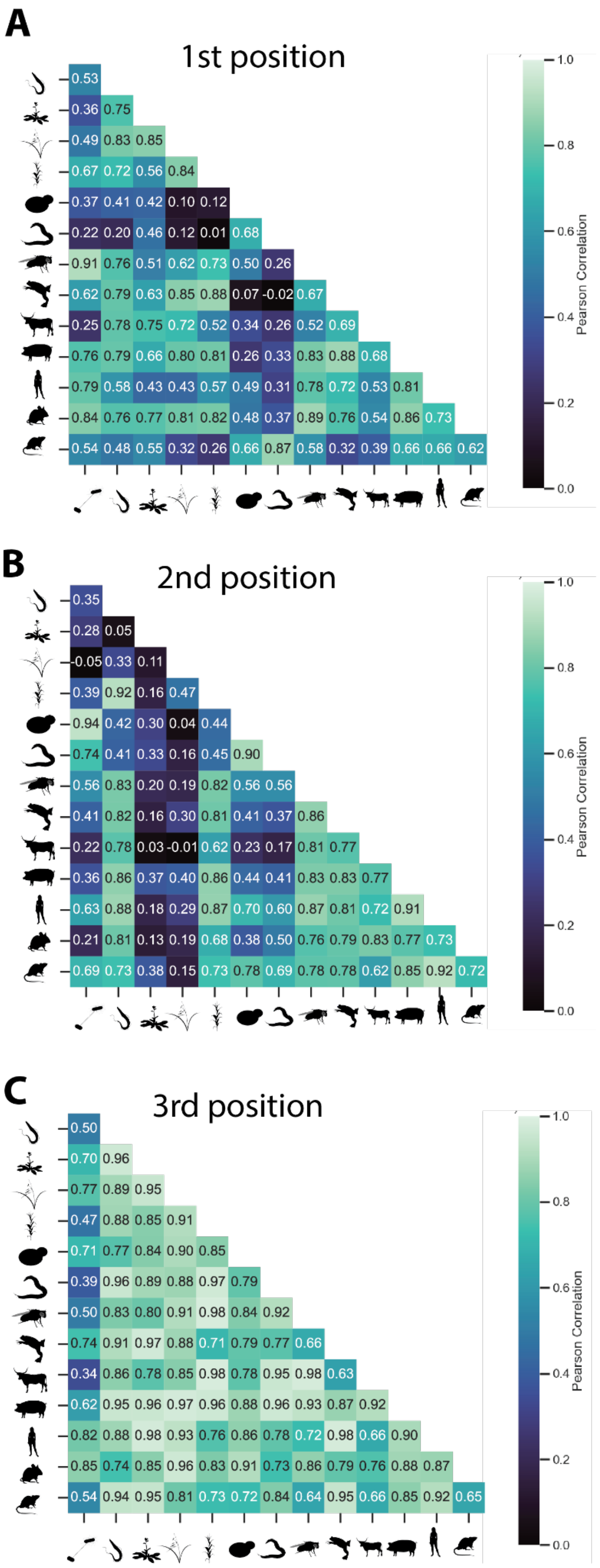


**Supplementary Figure 11: Correlation of nucleotide mismatches for all species pairs.** Cells indicate Pearson's r between nucleotide mismatch proportions (see Suppl. Fig. 10) for first (A), second (B), and third (C) codon position.

**Supplementary Table 1. Data overview.**

| Species | Datasets | Detected proteins | Proteins with misincs. | % Proteome | % Proteome with misincs. | Total spectra identified [mio.] | Misinc. spectra | Misinc. positions | Log10 misinc. rate |
|---|---|---|---|---|---|---|---|---|---|
| *E. coli* | 100 | 3,831 | 1,177 | 85.5 | 30.7 | 759.4 | 72,076 | 11,062 | -4.046 |
| *T. brucei* | 42 | 8,561 | 943 | 81.9 | 11.0 | 152.3 | 13,858 | 1,837 | -4.046 |
| *A. thaliana* | 156 | 27,498 | 2,085 | 66.7 | 7.6 | 1,389.0 | 38,925 | 7,917 | -4.523 |
| *O. sativa* | 17 | 43,673 | 686 | 27.0 | 1.6 | 177.6 | 5,893 | 1,380 | -4.523 |
| *Z. mays* | 17 | 39,208 | 698 | 32.5 | 1.8 | 130.6 | 5,676 | 1,008 | -4.398 |
| *S. cerevisiae* | 100 | 5,888 | 1,182 | 91.9 | 20.1 | 742.8 | 21,699 | 5,876 | -4.523 |
| *C. elegans* | 43 | 19,838 | 705 | 50.0 | 3.6 | 413.1 | 6,841 | 2,283 | -4.699 |
| *D. melanogaster* | 62 | 13,821 | 1,622 | 78.0 | 11.7 | 371.2 | 21,294 | 4,145 | -4.222 |
| *X. laevis* | 26 | 34,806 | 1,013 | 37.9 | 2.9 | 211.5 | 22,203 | 2,005 | -3.959 |
| *B. taurus* | 29 | 23,844 | 618 | 50.6 | 2.6 | 139.5 | 8,639 | 1,578 | -4.222 |
| *S. scrofa* | 34 | 22,803 | 1,114 | 55.5 | 4.9 | 252.7 | 17,003 | 2,761 | -4.155 |
| *H. sapiens* | 1,908 | 20,594 | 5,776 | 91.1 | 28.1 | 11,231.2 | 421,922 | 44,661 | -4.398 |
| *M. musculus* | 368 | 21,968 | 2,298 | 79.9 | 10.5 | 2,283.2 | 48,973 | 9,737 | -4.699 |
| *R. norvegicus* | 116 | 22,816 | 1,227 | 66.9 | 5.4 | 1,080.6 | 25,979 | 3,974 | -4.699 |

**Supplementary Table 2: Undetectable misincorporation types.** Listed are the misincorporations types that cannot be detected with our pipeline, because the mass shift is very similar to a post-translational modification or artifact of sample preparation.

| **Origin residue** | **Destination residue** | **Delta Mass [u]** | **Masking PTM** |
|---|---|---|---|
| S | D | 27.99491 | Formylation |
| D | E | 14.01565 | Methylation |
| T | E | 27.99491 | Formylation |
| S | T | 14.01565 | Methylation |
| F | Y | 15.99491 | Oxidation |
| N | C | 45.98772 | Beta-methylthiolation |
| N | Q | 14.01565 | Methylation |
| Q | E | 0.98402 | Deamidation |
| S | E | 42.01056 | Acetylation |
| P | E | 31.98983 | Dioxidation |
| N | D | 0.98402 | Deamidation |
